# Neural dynamics reveal foraging-like computations in the frontal cortex

**DOI:** 10.1101/2025.09.29.679289

**Authors:** Zsombor Ungvárszki, Clément Goussi-Denjean, Anna Székely, Gergo Orbán, Matteo di Volo, Emmanuel Procyk

**Affiliations:** Université Lyon 1, INSERM, Stem Cell and Brain Research Institute U1208, 69500 Bron, France; Department of Computational Sciences, HUN-REN Wigner Research Center for Physics, Budapest, Hungary; Department of Cognitive Science, Faculty of Natural Sciences, Budapest University of Technology and Economics, Budapest, Hungary

**Keywords:** Decision making, Computational modeling, Strategy switching, Macaque electrophysiology, Population coding

## Abstract

The prefrontal cortex has a central role in flexible behavior. Computations for weighing alternatives, committing to an option, integrating evidence, and abandoning an option are dominantly formalized in reinforcement learning (RL). Yet, ecological considerations showcase foraging as a competing account. Identifying how the brain actually solves efficient weighing of options requires the identification of the neural representation of this abstract yet precise computation. We recorded neural populations from three areas of the primate frontal cortex involved in decision making to characterize neural dynamics while animals performed a standard RL task. By combining behavioral and neuron population analysis, we found a neural subspace in midcingulate (MCC) and ventrolateral prefrontal cortex (vLPFC) that precisely tracked value computations. We demonstrated that MCC dynamics showed signatures of foraging-like computation but conflicted with RL requirements. Our study thus provides a detailed neural account of foraging-like flexibility in the primate frontal cortex.

## Introduction

To thrive in our natural environment, alternative options need to be assessed in order to find the one that is worth investing in. A formal understanding of this challenge is critical for understanding prefrontal cortical functioning in healthy and pathological conditions. To address this, value guided choice has been extensively investigated (Rangel et al., 2008; Haber and Knutson, 2010) and multiple computational frameworks have been put forward: learning theory offers reinforcement learning as a general framework to learn from action outcomes (Sutton, 2018); in parallel, ecological motivations highlighted the importance of a foraging account, a more limited but effective computational solution in naturalistic reward seeking behaviors (Charnov, 1976; Pearson et al., 2014; Alejandro and Holroyd, 2024). Reinforcement learning instructs to calculate the value of all alternatives and choose the one with the highest value. In contrast, instead of explicitly comparing the value of alternatives, the foraging account of choice only assesses the option that is currently viewed the best by calculating the value of persisting or abandoning it (Charnov, 1976). While behavioral models can find a quantitative advantage of one account over the other when predicting choices (Constantino and Daw, 2015; Alejandro and Holroyd, 2024; Zid et al., 2025), these differences are subtle compared to the qualitative differences expected in the neural representations that serve the computations. Hence, the critical question is whether it is the foraging or the RL framework that neural dynamics support.

Multiple prefrontal cortical subregions, striatal loops, and aminergic systems contribute to value guided choices (Montague et al., 2004; Aston-Jones and Cohen, 2005; Haber and Knutson, 2010). In primates, a part of the frontal cortical network, the midcingulate cortex (MCC), was shown to integrate multiple factors affecting the value of a choice and to encode action or task values (Amiez et al., 2006; Quilodran et al., 2008; Kennerley et al., 2011; Kolling et al., 2012; Khamassi et al., 2015; Hunt et al., 2018; Muller et al., 2024). MCC is involved in the integration of experience across time, as shown by its sensitivity to the history of past outcomes (Kennerley et al., 2006; Seo and Lee, 2007; Kennerley et al., 2011; Wittmann et al., 2016), as well as by a weighing of outcomes commensurate with the volatility of the environment (Behrens et al., 2007). Hence, the MCC is endowed with the ingredients necessary for evaluating, updating, and maintaining behavioral strategies in tasks requiring flexibility like in foraging (Hayden et al., 2011; Holroyd and Yeung, 2012; Kolling et al., 2016).

Subjective values and monitoring in MCC have been associated with high level adaptive functions, supported by larger frontal networks, and contributing to exploratory decisions, shifting and information seeking (Stoll et al., 2016; Tervo et al., 2021; Monosov, 2024). For instance, MCC might contribute to adaptive regulations of cognitive control and planning in interactions with the dorsolateral prefrontal cortex (dLPFC) (Shenhav et al., 2016; Kolling et al., 2018; Gandaux et al., 2025). Further, through its interactions with the ventro-lateral prefrontal cortex (vLPFC), MCC might serve efficient adaptation by linking information seeking and credit assignment (Monosov and Rushworth, 2022). Interestingly, the different intrinsic neural timescales and dynamics observed in these frontal areas suggest different computational capacities and a putative dissociation in the mechanisms enabling information integration (Fontanier et al., 2022; Procyk et al., 2021). However, whether values integrated in some of these areas are built thanks to action selective processes or foraging-like computations remains to be answered. Resolving this question requires the identification of the actual value updating mechanism in neural activity. Dissecting how behaviorally relevant variables are multiplexed or orthogonalized in different neural subspaces, and how they are dynamically evolving during task epochs provides the opportunity to unveil how neural networks represent associations or organize complex information (Stoll and Rudebeck, 2024; Chen et al., 2024). A major goal here is thus to decipher from frontal cortex neural dynamics whether flexible behavior relies on RL-like or foraging-like updating of subjective values.

In this study, we tracked value computations in neural population responses across three prefrontal cortical subregions (MCC, dLPFC and vLPFC) while animals needed to deliberate multiple options for obtaining reward. The probabilistic three-armed bandit task warranted the use of an RL strategy but could also be solved with a foraging approach. Central to both frameworks was the calculation of a value variable, but it had divergent roles as RL associates value to alternative options, while foraging associates it to whether the current option is to be abandoned. This provided us an opportunity to identify signatures specific to the alternative frameworks in neural representations. Using behavioral modeling, we inferred the value of the animal’s choice, which precisely integrated the outcome of past choices. Our analysis revealed that the choices of the experimental animals adhered to optimal integration of outcome history. Alternative models could be contrasted solely based on behavior, which revealed similar predictive power of alternatives with a marginal advantage of the foraging account. By capitalizing on the fact that the value derived from behavior provided a trial-by-trial measure of the subjective value of the options the animals had in a trial, we used it to regress neural recordings. We showed differential neural coding in the three subregions, with MCC and vLPFC representing value but not dLPFC. Importantly, by parametrically tuning the computational model of behavior, we have shown that the behaviorally derived value was optimal to fit the neural data. Critically, our analyses demonstrated that the neural value identified in MCC carries the signature of foraging-like value computations and contradicts the prediction of reinforcement learning-based value computations. The value computations implied by the foraging model permitted us to track the dynamics of value updating across different time points in a trial, on a trial-by-trial basis in the MCC and vLPFC. Dimensionality reduction and trajectories in neural state spaces revealed a functional dissociation between regions, as precise temporal unfolding of value updating could be identified in MCC but not in the other recorded prefrontal areas.

## Results

We trained two macaque monkeys on a three-armed bandit task in which the payoff was stochastic and stable in blocks of trials (**Fig. 1A-C**). In each block one potential target had a high (70%) while the two others had a low (25%) reward rate (**Fig. 1B**). Reward rates remained stable throughout blocks of 40*±*5 trials, but were randomly shuffled between blocks (**Fig. 1C**). Optimal behavior required choosing the target with the high reward rate and continuing to choose it as long as its payoff remained high, and switching to a new target only when its payoff decreased. Since rewards were not delivered deterministically, a single unrewarded trial could not identify a switch in payoffs, but animals had to integrate the successive trial outcomes to identify a drop in reward rate. Block ends were not signaled to the monkeys, thus efficient performance required constant evaluation of the selected option to identify a change in the environment. Monkeys chose the optimal target in an average of 72% of trials (KA: 78*±*6% across sessions, PO: 64*±*12%), performing well above the level of random choice (33%), systematically exploring alternatives after block changes and settling on one target later in the block (**Fig. 1D-E**). The performance of the animals was close to an optimal model (81%; see Methods section), demonstrating that monkeys understood the task structure well and developed an efficient strategy. Importantly, switching after a positive outcome was almost absent for KA (1% of all positive outcomes) and was rare for PO (6%), which indicates that animals followed a strategy that exploits the current option if rewarded, and explores new options mostly in light of unrewarded trials. Following a negative outcome, animals switched on average around 30% (KA: 23%; PO: 40%) of trials, indicating that they do not simply follow a win-stay-lose-switch strategy. Instead, the recent outcome history influenced switching probability. Logistic regression revealed that the outcome of the last six trials significantly influenced switching decisions in both monkeys (**Fig. 1F**; prediction accuracy for KA: 84%; PO: 77%). Regression weights followed an exponential recency-weighting (**Fig. 1F**), indicating that monkeys precisely integrated reward history over multiple trials, in line with a standard prediction-error learning rule (Sutton, 2018). Animals identified the high payoff target in around 10-15 trials after a block boundary, and reliably exploited this target until the upcoming block boundary (**Fig. 1E**).

**Figure 1.**
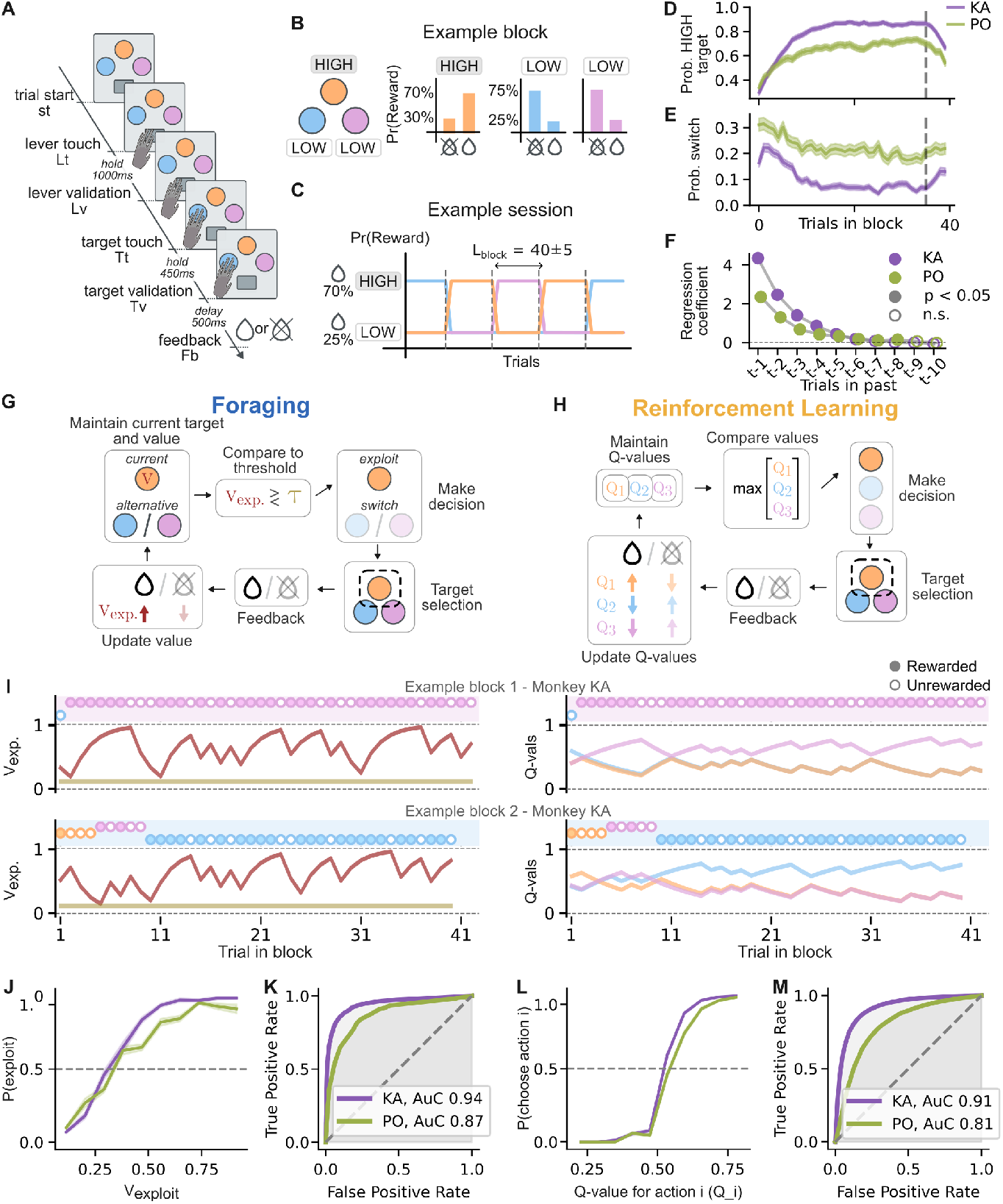
Behavioral performance and modeling. **A-C**, Behavioral paradigm. **A**, In each trial, monkeys first had to initiate the trial by touching a lever and holding it for 1000 ms (lever touch to validation, Lt, Lv, respectively), then choose a target by holding it for 450 ms (target touch to target validation, Tt, Tv, respectively). After a 500 ms delay, either a juice reward or no reward was delivered (Fb), depending on the trial outcome. B, Outcome probabilities of a trial: in each trial, one target had a 70% reward rate and the other two had 25% reward rates (HIGH and LOW, respectively). **C**, The identity of the HIGH target changed randomly after blocks of 40±5 trials. **D**,**E**, Percent HIGH target selection (D) and probability of switching (E) throughout a block, across sessions for the two monkeys (KA, PO). Dashed line marks the start of the last five trials, where reward probabilities change gradually. **F**, Logistic regression of the contribution of past trials to choosing to stay with the previous target or switching to a new one from most recent (t-1) to a distant (t-10) trial, for the two animals. Filled circles indicate significant deviation from zero. **G**,**H**, Schematic illustration of the computations for choice foraging and reinforcement learning, respectively. The Foraging model tracks the value of the currently pursued action (*V*_exploit_) and compares it to a threshold (*τ*) to decide to persist or switch to an alternative action. The reinforcement learning (RL) model tracks action values – one per option – and compares them directly to select the best one. **I**, Choices in two example blocks and calculated values for the two theoretical accounts. Left, the Foraging model maintains a single running value (red); as it approaches the switching threshold (light brown line), the probability of switching increases. Right, the Reinforcement Learning model tracks three action values parallelly and selects the highest. **J**,**K**, Characteristics of the Foraging model. **J**, Probability of persisting (exploit) or switching as a function of the exploit value (*V*_exploit_). **K**, Predictive accuracy of (*V*_exploit_) quantified by ROC-AUC. **L**,**M**, Characteristics of the RL model. **L**, Choice probability as a function of the Q-value (probabilities computed per choice then averaged). **M**, Predictive accuracy of Q-values quantified by ROC-AUC (averaged across three Q-values).

### Normative models of behavior

To precisely characterize behavior, we designed quantitative models that we fitted to the choices of the animals on a trial-by-trial basis. We considered two main alternatives according to the key theoretical accounts proposed for value-guided choice. First, we set up a foraging model, which is constrained to evaluate the viability of the current choice but not the alternatives. This foraging model calculates the choice value for the target actually visited but does not represent the value of specific alternatives (Charnov, 1976; Zid et al., 2025). Second, reinforcement learning (RL) was recruited to assess the value of choice, which directly evaluates all potential alternative choices by assigning values for all of these (Sutton, 2018).

The foraging model implements a simple compare-to-threshold decision: exploit the current option if its reward rate is above the average reward rate in the environment, and switch otherwise (Charnov, 1976; Zid et al., 2025). (**Fig. 1G**, see Methods for details). The value of exploitation is estimated trial-by-trial according to a standard prediction-error framework using the delta learning rule (Sutton, 2018) (see an example session in **Fig. 1I**). This value is then compared to a fixed threshold to decide to stay with the currently selected target or switch to an alternative. In general, the threshold is determined by the expected global payoff in the task, but we considered this threshold as a free parameter. Assessment of value is performed after each feedback, and its update depends on a single parameter, the learning rate. Intuitively, this learning rate determines the relative importance of the actual value estimate and the current feedback. Consequently, the learning rate directly affects the rate at which the influence of an earlier feedback is discounted, resulting in an exponential depreciation of earlier feedback (Sutton, 2018) (**Supplementary Fig. S1**).

For an RL account of value-guided choice, we used a Q-learning model, which implements a compare-to-alternatives decision (Sutton, 2018). The RL model assigns values to each action and compares these alternative options to reach a decision (**Fig. 1H**). We set up two flavors of the RL model. First, a standard Q-learning model was used, which consistently updates the value of the current choice upon new information obtained through feedback, but the value estimates of alternatives are assumed to be independent and are consequently not affected by feedback on the current choice. We also implemented an inferential RL model that is capable of learning about the dependencies of the payoff of alternatives. This model is reminiscent of the ideal observer model that maintains hidden variables associated with the reward rates of alternative targets and makes inferences about which of the alternatives is the one characterized by the high payoff. Action values are updated based on the delta learning rule: for the selected target, the prediction error is given as the difference between its previous value and the observed outcome. For the unchosen options, the model uses the inverse of the outcome to compute a prediction error. Formally, value updates are similar to those of the foraging model, as a learning rate parameter determines the relative importance of the current action value and current feedback. However, in contrast with the foraging model, polarity of action value updates depends on what choice the actual feedback belonged to (**Fig. 1I**; Methods; **Fig. 2A,B**). Action selection was implemented with a softmax rule, and to give the RL model more flexibility, we added an additional *stickiness* parameter that biases repeating the previous action, consistent with the task structure which often rewards repeating actions.

**Figure 2.**
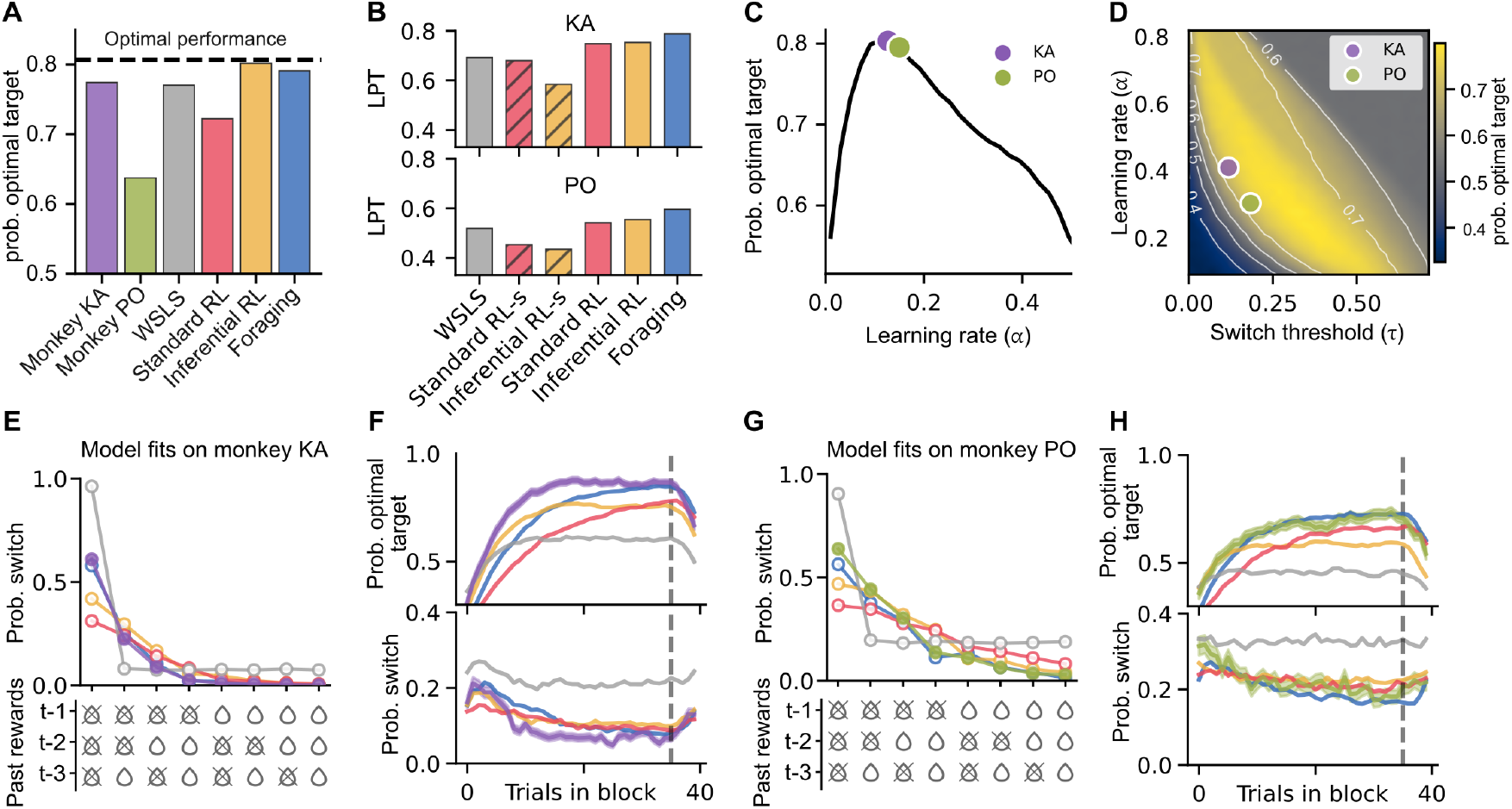
Performance of alternative models. **A**, Efficiency of models and experimental animals in harvesting rewards. Four models are compared: win-stay–lose-switch (WSLS), two versions of the reinforcement learning model (Standard RL, Inferential RL), and the foraging model. Chance level is at 33%. Dashed line: performance of an optimal model. **B**, Predictive performance of alternative models on the choice behavior of the two animals. Both RL models were tested with and without fitting the stickiness parameter (RL-s). Performance is measured through calculating the likelihood of making the choice of the animal per trial. **C**, Comparison of Inferential RL performance at various values of learning rate and fitted values for the two animals (dots). **D**, Comparison of foraging model performance at various settings of two key parameters (learning rate and switch threshold) and the performance of the model at parameter values fitted to the choices of the experimental animals. **E**,**G**, Behavior of the alternative models (colors and labels as on A) for different reward histories, evaluated through the probability of choosing to switch in trial *t* (E: KA; G: PO). **F**,**H**, Model predictions of the alternative models for within-block behavior. Models are fit to individual animals (F: KA; H: PO). Behavior is best explained by the foraging model for both monkeys. Top: probability of selecting the HIGH target; bottom: probability of choosing to switch between trials. Dashed line marks the start of the last five trials, where reward probabilities change gradually

Additionally, as a simple baseline that does not track values and prediction errors, we implemented a version of the win-stay-lose-switch model (WSLS) which switches if the last three outcomes were all unrewarded and stays with the previously selected target otherwise. This model reflects our observation that both monkeys switched with at least 50% probability only after experiencing at least three negative outcomes (**Fig. 2E,G**).

Crucially, both the foraging and the RL models can predict animals’ choices, despite predicting different decision variables: the foraging model predicts switching from a single value of exploitation, whereas the RL model predicts actions from three action values. When fit to behavior, the value of exploitation accurately predicted switches and the action values accurately predicted actions (**Fig. 1J,L**), as reflected by receiver operating curve analysis (ROC) (**Fig. 1K,M**), where area-under-curve values were high for both animals. However, these analyses do not permit a direct comparison between models, so we next compared them explicitly.

To check if models can, in principle, solve the task efficiently, we optimized models for reward maximization. In this validation, the inferential RL model slightly but significantly outperformed the foraging model (percent optimal target selection: foraging: 79%, inferential RL: 80%), though both models performed just below a normative benchmark (81%; see Methods) (**Fig. 2A**). Both of these models outperformed the standard RL model (72%) and the win-stay-lose-switch model (77%). All model performances differed significantly from each other (see **Supplementary Table S1** for pairwise comparisons). Similar performance of the foraging and RL models was guaranteed by sharing the delta learning rule, resulting in correlated updates of value estimates.

To assess whether these models explain the choices of monkeys, we fit the models to the behavior using the log-likelihood method. The foraging model showed better predictive power than both the inferential RL model and the standard RL model across the two animals (likelihood per trial (LPT): KA – foraging: 0.79, inferential RL: 0.75, standard RL: 0.75; PO – foraging: 0.60, inferential RL: 0.55, standard RL: 0.54) (**Fig. 2B**). Importantly, without the stickiness parameter, both RL models fit substantially worse (**Fig. 2B**, RL-s represents models without the stickiness parameter). Moreover, the foraging model consistently outperformed the WSLS model (WSLS LPT: KA: 0.69; PO: 0.52) (**Fig. 2B**).

The fitted parameters of the inferential RL and the foraging models largely overlapped with those that maximize expected rewards across different parameterizations (excluding the noise parameter in both models and the stickiness parameter as well in the RL model; **Fig. 2C,D** respectively). This suggests that while both models capture the structure of optimal choices, they differ in accounting for systematic biases – together with its higher LPT, the foraging model appears to explain these biases more accurately.

To assess where the fitted models reproduce – and critically, differ from – the monkeys’ strategy, we simulated extended sessions (100,000 trials) using parameters estimated for each animal. The foraging and both RL models closely matched the animals’ performance (**Fig. 2E-H**; percent optimal target selection: KA – animal: 78%, foraging model: 71%, inferential RL: 69%, standard RL: 62%; PO – animal: 64%, foraging model: 64%, inferential RL: 56%, standard RL: 57%), while the WSLS model performed substantially worse (KA: 58%; PO: 45%). The fitted models reproduced key characteristics of monkeys’ behavioral strategy: switching after rewarded trials was rare for both monkeys, while after unrewarded trials, the switching probability increased with the number of recent unrewarded outcomes (**Fig. 2E,G**). Furthermore, all models - except the baseline WSLS – learned to find the high-payoff target during a new block by progressively abandoning low-payoff targets until eventually settling on the best target (**Fig. 2F**).

### Single-unit activity reflects task variables

We recorded single-unit activity simultaneously from the midcingulate cortex (MCC) and the lateral prefrontal cortex (LPFC) from the two macaque monkeys (KA: **Fig. 3A**, PO: **Supplementary Fig. S2B**). Two linear 16-contact probes were driven perpendicular to the cortex acutely in each recording session, changing probe location systematically between sessions, thus allowing us to sample different neuron populations across sessions. Recordings covered a broad extent of MCC and LPFC (**Fig. 3A**). MCC recordings were in the fundus and dorsal bank of the cingulate sulcus spanning rostro-caudal levels where outcome and value-related coding have been reported, and encompassing multiple subregions (Procyk et al., 2016; Ducret et al., 2024). Recordings in the LPFC were in both the dorsal (dLPFC) and ventral (vLPFC) gyri around the principal sulcus (**Fig. 3A**). In total, we isolated 4,963 units (monkey KA: MCC: 1466; dLPFC: 964; vLPFC: 244; monkey PO: MCC: 1355; dLPFC: 794; vLPFC: 140). On average, we recorded 24*±*8 MCC, 20*±*9 dLPFC and 20*±*7 vLPFC units per session in monkey KA, and 26*±*9 MCC, and 17*±*9 dLPFC and 20*±*9 vLPFC units per session in monkey PO from all recording sessions (KA: MCC: 60; dLPFC: 48; vLPFC: 12; PO: MCC: 53; dLPFC: 46; vLPFC: 7) Activity of single neurons was investigated for the coding of key task variables such as feedback (outcome valence) and selected target. Further, we capitalized on recording the choices of animals to establish estimates of the value both in the foraging and RL models (see Methods). These behaviorally derived estimates were then used as regressors to the neural data. Single neuron activity displayed sensitivity to feedback, value, and target identity with heterogeneous coding dynamics (**Fig. 3B**).

**Figure 3.**
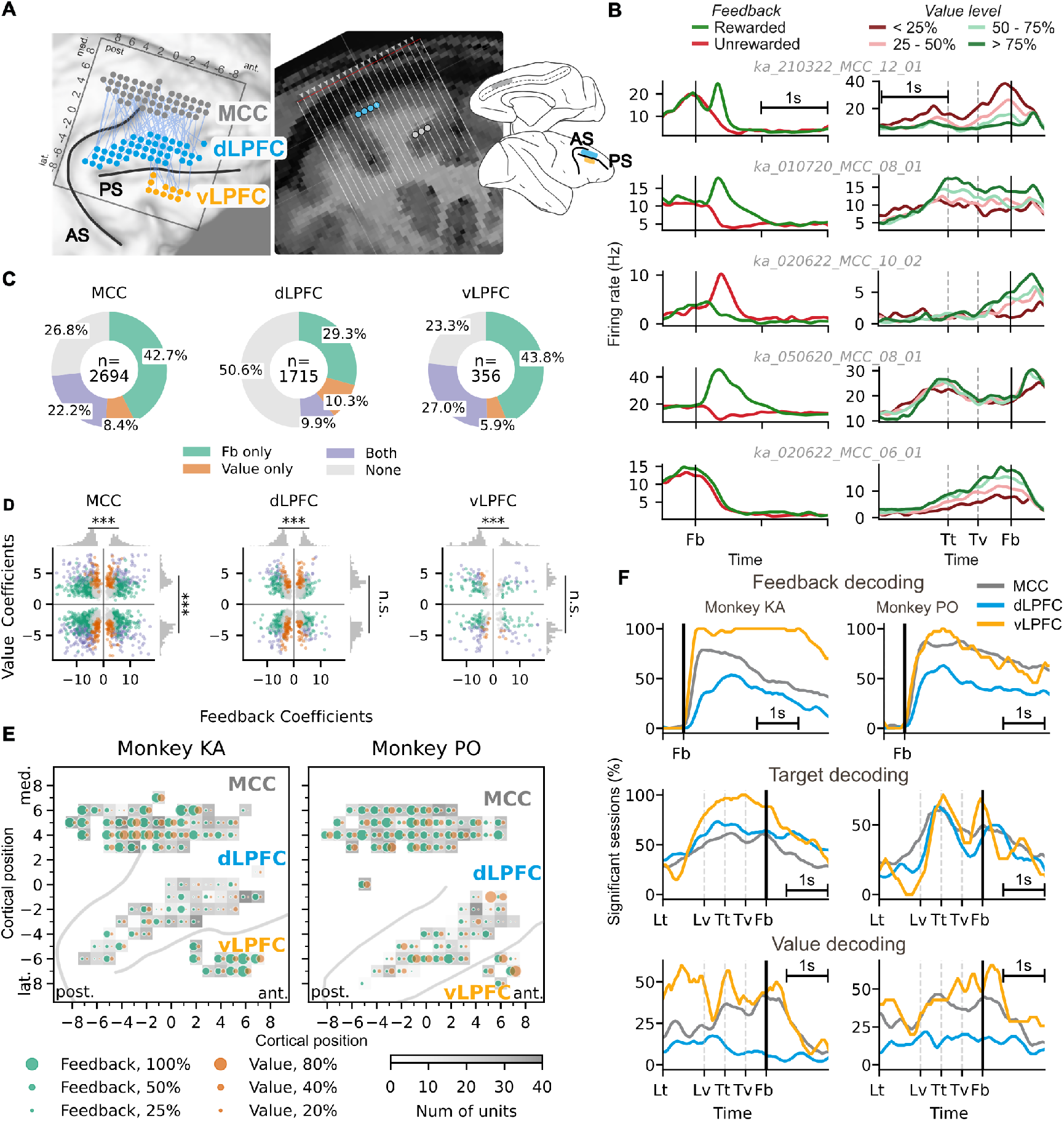
Representation of task variables in MCC and dLPFC. **A**, Electrode placements in the midcingulate cortex (MCC) and the dorsolateral and ventrolateral prefrontal cortex (dLPFC and vLPFC, respectively) of monkey KA. Recording site positions changing from session to session are marked with dots on the cortical surface (left) and cross sections (right). **B**, Time course of single unit firing patterns of five example units in MCC across a trial. Mean firing rates are shown for different trial outcomes (left column) and different levels of the behavior-derived value (right column). Values are derived from the foraging model. **C**, Fraction of units encoding feedback and value in the recorded regions. **D**, Level of contribution of feedback and value to the responses of individual neurons (dots) as measured by GLM coefficients. Units recorded from different regions are shown on different panels. Marginals (grey) show the distribution of GLM coefficients for individual variables. The numerosity of units with different polarity GLM weights is contrasted with binomial test. **E**, Distribution of units with significant feedback- and value-related CPD across the recording sites. Dot size indicates the proportion of units with contributions from the two variables. **F**, Temporal dynamics of encoding task variables in the three recorded brain regions, as reflected by the fraction of sessions with significant cross-validated linear decoder performance trained for trial outcome (top), chosen option (middle), and value (calculated using the foraging model based on choices, bottom).

To quantify neural encoding of task variables at the single-unit level, we applied a generalized linear model (GLM) to predict spike counts in sliding time windows (50 ms, Methods) based on two task variables: Feedback (outcome: 0/1) and Value (0-1) (see examples in **Fig. 3B**). Because these variables could be correlated, we computed the Coefficient of Partial Determination (CPD) for each, to characterize their unique contribution to the variance in the neural activity. The relative coding of variables varied between areas: more neurons in the MCC and vLPFC coded for feedback and/or value than in the dLPFC (MCC: 73.2%, vLPFC: 76.7%, and dLPFC: 49.4%) (**Fig. 3C,E**). Within each area, we found subpopulations encoding feedback only, value only, or both (example neurons of each case are shown in **Fig. 3B**), revealing widespread co-encoding of feedback and value, with many neurons representing both variables. Furthermore, single unit coding patterns were varied revealing neural activations for both positive (‘reward’) and negative (‘no reward’) feedback and both positive or negative activity modulation with value. Note, however, that significantly more units were more active for negative than for positive feedback in all areas (MCC: 1117 out of 1830 feedback-coding units; dLPFC: 419 out of 688; vLPFC: 177 out of 277; binomial test against chance level of 0.5, two-sided, *p <* 0.001 for all areas), consistent with previous findings (Quilodran et al., 2008; Matsumoto et al., 2007). Feedback and value were heterogeneously represented across MCC and LPFC sites with more frequent occurrence in the central part of the covered MCC region, although with interindividual variability, corroborating previous work (Procyk et al., 2016). Correlates of these variables were sparse in dLPFC (**Fig. 3E**). Due to limits of the recording chamber, we covered only a limited number of sites in a vLPFC subdivision (mostly area 46vc) but those revealed clear resemblance to MCC (see also further decoding analyses below).

We used population decoders to uncover the temporal dynamics of feedback and value encoding (using 50 ms sliding windows). Sessions were classified as coding for a variable if they showed significant decodability (*p <* 0.05) in at least four consecutive time bins. Value signals were steadily maintained during the inter-trial period in a small subset of sessions (**Fig. 3F**), but the number of value coding sessions ramped up towards the feedback time. Feedback became significantly decodable in all recorded areas in the majority of the sessions (maximum fraction of significantly decodable sessions: KA – MCC: 78.7%, dLPFC: 53.5%, vLPFC: 100%; PO – MCC: 87.6%, dLPFC: 63.1%, vLPFC: 100%; **Fig. 3F**), and remained decodable – though in fewer sessions – in the inter-trial period. Correlates of the chosen target could be identified in all areas (**Fig. 3F**), with decoding performance peaking at target touch and maintaining until feedback onset. Coding dynamics were consistent across animals (see **Supplementary Fig. S3A-C**). Note, however, that time on task effects (Goussi-Denjean et al., 2023) and the block structure of the task artificially contributed to the decoding performance.

In conclusion, descriptive analyses revealed distributed representations of outcome and value across the frontal cortex, with clear dynamics developing between feedback onset, target selection, and the next feedback. Representation of these variables in the population responses provides the opportunity to investigate the precise arithmetic of value-based choice in alternative models as well as to investigate the dynamics of value updates with experience.

### Precise arithmetic of outcome integration in the MCC

Decisions rely on the expected value of actions. In stochastic environments, this value must be inferred from experience. Calculation of value requires the integration of feedback history such that earlier experiences are precisely discounted. Close to optimal behavior identified by behavioral analysis indicates that monkeys are capable of delicately evaluating odds and outcomes. We therefore set out to investigate if the precise arithmetic of value inference from experience can be identified in neural responses. To characterize the dynamics of neural representation of outcome information across trials, we combined all recording sessions into a single composite session by aligning firing rates of single neurons across trials with identical outcome history (limiting the horizon to three trials into the past, see Methods for details).

We found that feedback information was steadily decodable (using logistic regression) in all areas after feedback delivery and during the inter-trial interval (ITI), persisting into the subsequent trial up until the next feedback delivery. After this point, decodability diminished rapidly (**Fig. 4A**). Note that as feedback history guides choices (**Fig. 1F**), the value signal carries information about the past feedback and therefore it can be decoded in subsequent trials. In those time windows, however, feedback from multiple trials is integrated and analyzed separately, in the coming paragraphs.

**Figure 4.**
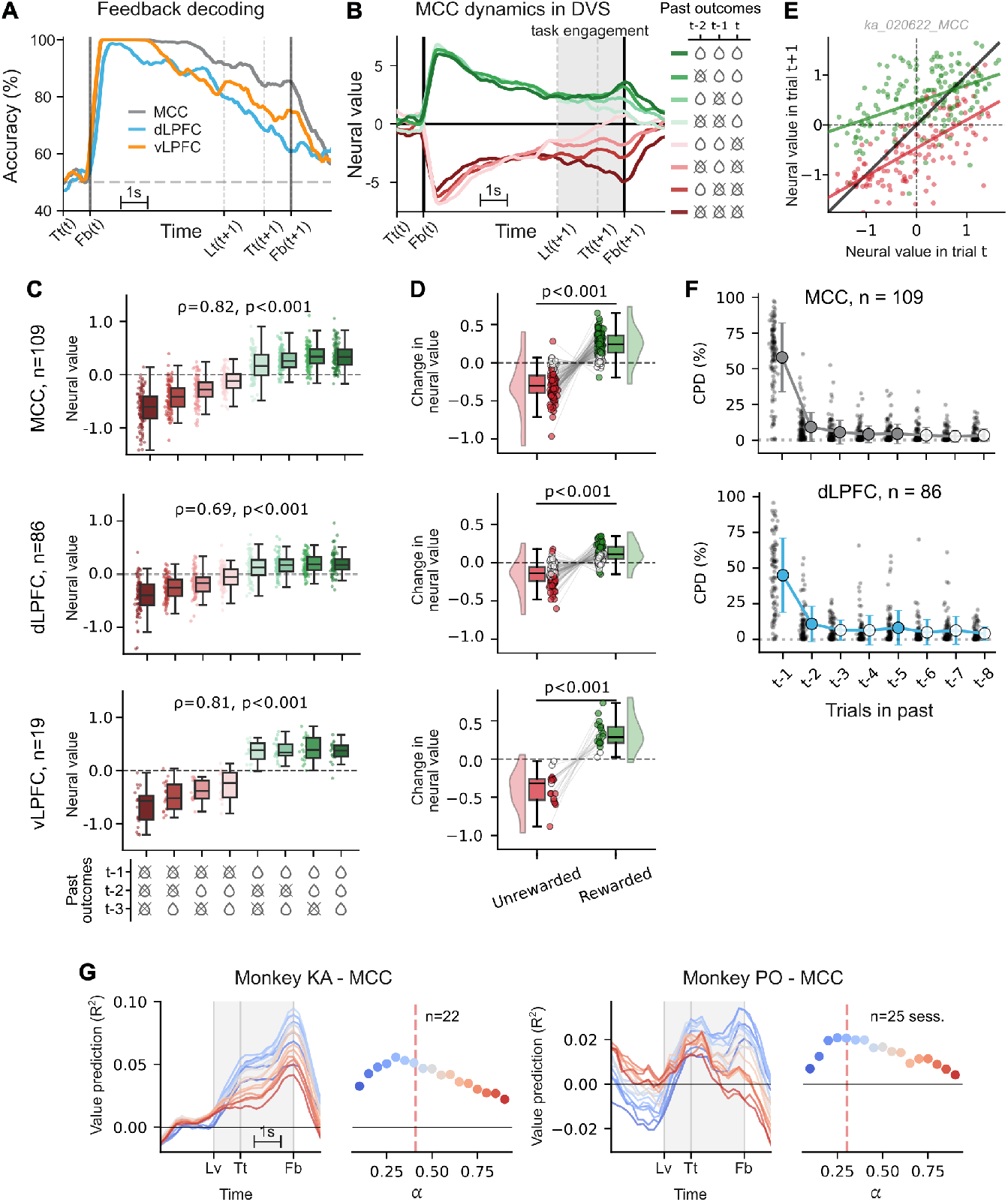
Arithmetic of neural dynamics. **A**, Time course of feedback decoding performance in the three recorded regions. Lines show mean decoder performance in the composite-session. **B**, Dependence of MCC activity on outcome history. Feedback decoders are trained for individual time points during a trial of the composite-session. Trajectories are population responses projected in the one-dimensional subspace identified by the decoder, average population response intensities are calculated for different outcome histories (colors). This quantity is called the *neural value*. Shaded background: period of task engagement. **C**, Across-session dependence of neural value on outcome history for the three recorded regions. Dots represent individual sessions, box and whisker shows median and (10, 25, 75, 90 percentiles), neural value is measured in the task engagement period, *n* indicates number of sessions. **D**, Trial-by-trial change of neural value across sessions for positive and negative outcomes, for the three recorded regions. The distribution of changes across sessions (dots) is characterized with violin, box-and-whisker plots. **E**, Trial-by-trial (dots) dynamics of the neural value in an example session from MCC. Positive outcome (green) raises the neural value more if it was low in the previous trials, while a negative outcome (red) decreases the neural value more if it was high in the previous trial. Neural value is calculated from the task engagement period. **F**, Contribution of past outcomes to the neural responses in MCC and dLPFC, as quantified by CPD. Dots indicate individual sessions, circles and error bars indicate mean and standard deviation. Filled circles indicate significant deviation from zero. **G**, Optimality of the behavior-derived value for predicting neural activity. Left panel: Correlation of calculated behavioral value with population responses as a function of assumed learning rates (*α*) of the foraging model (colors correspond to learning rates on the right panel) at different times during the trial. Shaded area: period of task engagement. Right panel: mean predictive strength of behavioral value as a function of learning rate, averaged across the task engagement period. Dashed line: learning rate from behavioral fit.

To analyze value computations in detail, we used a one-dimensional subspace of the neuron population activity in which the evolution of value can be tracked. For this, we used the subspace defined by the vector along which the signal associated with the value is strongest, the decision vector subspace (DVS) (Hajnal et al., 2023). We identified DVS for every time point during the trial and projected neural activity into this subspace. We refer to the level of population activity in DVS as the *neural value* signal. DVS revealed a systematic modulation of neuron population activity by recent outcome history, with the strongest modulation observed during the task-engaged period of the upcoming trial. This effect was strongest in MCC (**Fig. 4B**), and weaker but present in both LPFC subregions (**Supplementary Fig. S4A,B**; see quantitative analysis below).

Graded representation of the value signal becomes evident in a session-by-session analysis. We analyzed the neural value in the period between lever touch and feedback (a 2-s window; grey zone in **Fig. 4B**). Statistical testing confirmed a significant modulation of the signal by outcome history at the population level for all areas (Spearman Rho-test, *p <* 0.001 all areas), with the strongest effect observed in MCC and vLPFC (Spearman Rho test: MCC: *ρ* = 0.82, *p* = 6*×*10^*−*213^; dLPFC: *ρ* = 0.69, *p* = 5*×*10^*−*100^; vLPFC: *ρ* = 0.81, *p* = 4*×*10^*−*36^ **Fig.4C**). Note that one can observe a higher range of modulation for trials with recent negative outcomes in all three regions (positive sequences: MCC: *ρ* = 0.268, *p* = 1.2*×*10^*−*8^; dLPFC: *ρ* = 0.117, *p* = 3.0*×*10^*−*2^; vLPFC: *ρ* = 0.005, *p* = 9.6*×*10^*−*1^; negative sequences: MCC: *ρ* = 0.582, *p* = 6.2*×*10^*−*41^; dLPFC: *ρ* = 0.458, *p* = 3.4*×*10^*−*19^; vLPFC: *ρ* = 0.473, *p* = 1.6*×*10^*−*5^). Furthermore, the neural value signal displayed trial-by-trial dynamics consistent with expectations: neural value increased reliably following positive outcomes and decreased following negative outcomes, reflecting that its updating depended on outcome valence (**Fig. 4D**). This effect was significant across all three areas (Mann–Whitney U test, two-sided: MCC: *U* = 191, *n*_1_ = *n*_2_ = 109, *p* = 5*×*10^*−*35^; dLPFC: *U* = 399, *n*_1_ = *n*_2_ = 86, *p* = 5*×*10^*−*24^; vLPFC: *U* = 0, *n*_1_ = *n*_2_ = 19, *p* = 1*×*10^*−*7^), indicating trial-by-trial accumulation of feedback information in this subspace.

Single-session analyses further revealed that the dynamics of neural value updating follow theoretical predictions of delta-rule: the updating of the neural value depends both on the outcome and the initial value. Specifically, the updated value, *V* (*t* + 1), exceeded the initial value, *V* (*t*), after positive outcomes when *V* (*t*) was low (large prediction error), but increased only modestly when *V* (*t*) was high (small prediction error). Conversely, negative outcomes produced strong decreases in *V* (*t* + 1) relative to *V* (*t*) when the initial value was high, but only slight decreases when the initial value was low (**Fig. 4E**). This pattern was robust across monkeys and areas. To test this formally, we fit a single general linear model on V(t+1) that included *V* (*t*), outcome, and their interaction. A session was labeled delta-consistent if: (i) dependence on *V* (*t*) was significant for both outcome conditions; (ii) both regression slopes lay between 0 and 1; and (iii) the intercept for positive outcomes exceeded that for negative outcomes. Under *p <* 0.05, 46.3% (99 of all 214 sessions) sessions passed all three criteria (KA – MCC: 41/58, 70.7%; dLPFC: 10/46, 21.7%; vLPFC: 2/12, 16.7%; PO – MCC: 28/51, 54.9%; dLPFC: 14/40, 35.0%; vLPFC: 3/7, 42.9%), suggesting that frontal neural networks contribute to implementing a delta learning rule to track outcomes over multiple trials.

The graded neural value signal in DVS indicates integration of outcomes with a horizon beyond the immediate feedback. To assess the precise contribution of past outcomes, we analyzed the effect of outcome recency. We fitted a generalized linear model (GLM) regressing the neural value signal against the outcomes from the previous eight trials. For each historical outcome, we computed the coefficient of partial determination (CPD) to quantify its unique contribution (**Fig. 4F**). CPDs were computed separately for each session and then averaged across sessions. The analysis revealed a recency-weighted integration, with more recent outcomes exerting a stronger effect on the neural signal, and a gradual decay towards zero (**Fig. 4F**). Crucially, this pattern closely mirrors our observation in the behavioral analysis (see **Fig. 1F**). The timescale of this effect varied slightly across regions: in the MCC, the last 5 outcomes contributed significantly (permutation test, *p <* 0.05), while in dLPFC, the last 3 outcomes were significant. Due to the low number of recording sessions in vLPFC, no sufficient statistical power was available to carry out this analysis.

Value based decision in an environment with stochastic outcomes requires the discounted integration of the history of outcomes. This can be identified both in behavioral analysis (**Fig. 1E**) and in the analysis of neural responses (**Fig. 4F**). A tight link between behavioral discounting and neural discounting of past outcomes requires a match between the rates of discounting. We investigated this mapping from behavior to neuron population activity by fitting neural activity with different learning rates (the learning rate determines the effective discounting rate; see **Supplementary Fig. S1**). If a tight link exists, the best link is expected to match the behaviorally derived learning rate. Fitting the neuron population activity with varying learning rates displayed a distinctive maximum – this peak was well aligned with the behaviorally derived learning rates for the two animals in the MCC (**Fig. 4G**). Note that in the vLPFC, the single most recent outcome (reflected in maximal learning rate *α*) explained neural activity well (**Supplementary Fig. S5**). In the dLPFC, too few sessions met the significance criterion (8 of 94, 8.5%; see Methods).

In conclusion, neural decoding in frontal areas, and especially in the MCC, revealed clear correlates of trial-by-trial value updating and outcome integration, suggesting the use of a delta learning rule. The remaining question is thus whether this fine neural arithmetic reflected the use of a foraging-inspired compare-to-threshold or RL-like ‘compare-to-alternatives’ strategy.

### Neural hallmarks of foraging strategy

Having established that the foraging model is only marginally better than the RL model at the behavioral level, we asked which strategy better explains the neural data. Because the two models implement fundamentally different internal mechanisms – and thus different latent representations drive behavior despite similar observed choices – we used single-unit recordings from the prefrontal cortex to test whether algorithmic-level signatures of foraging versus reinforcement-learning are reflected in neural activity. We capitalized on the insight that the foraging model focuses on evaluating the value associated with the actual target, while the RL models maintain distinct values for alternatives. Specifically, a standard RL model compares the actual value to the last value of alternatives, while an inferential RL model continuously refreshes the values of alternatives. We test this crucial difference in the neural data by analyzing the neural subspaces identified by the value decoders. The difference between the foraging and RL frameworks is evident upon switching between targets: the neural value subspace identified for one target is expected to be unchanged when the target is switched for the foraging model. Thus, changes observed for the previous target upon receiving feedback are expected to show the same polarity for the new target (**Fig. 5A**). However, the updating of the target also updates the neural value subspace for RL models. For a standard Q-learning model the initial subspace will not be affected by feedback for the new target. For the inferential RL model, a more drastic distinction is expected, since a positive outcome for an alternative target decreases the value of the previous target. To test if the prediction of the foraging model or those of the RL models hold, we grouped trials based on the selected target and trained a linear decoder of model-based value estimates using trials from one target group. We then projected all neural activity in the subspace defined by this decoder. To obtain a trial-by-trial measure of value representation, we averaged this one-dimensional projection in the 2-second window spanning from lever touch (response onset) to feedback, resulting in a single value per trial.

**Figure 5.**
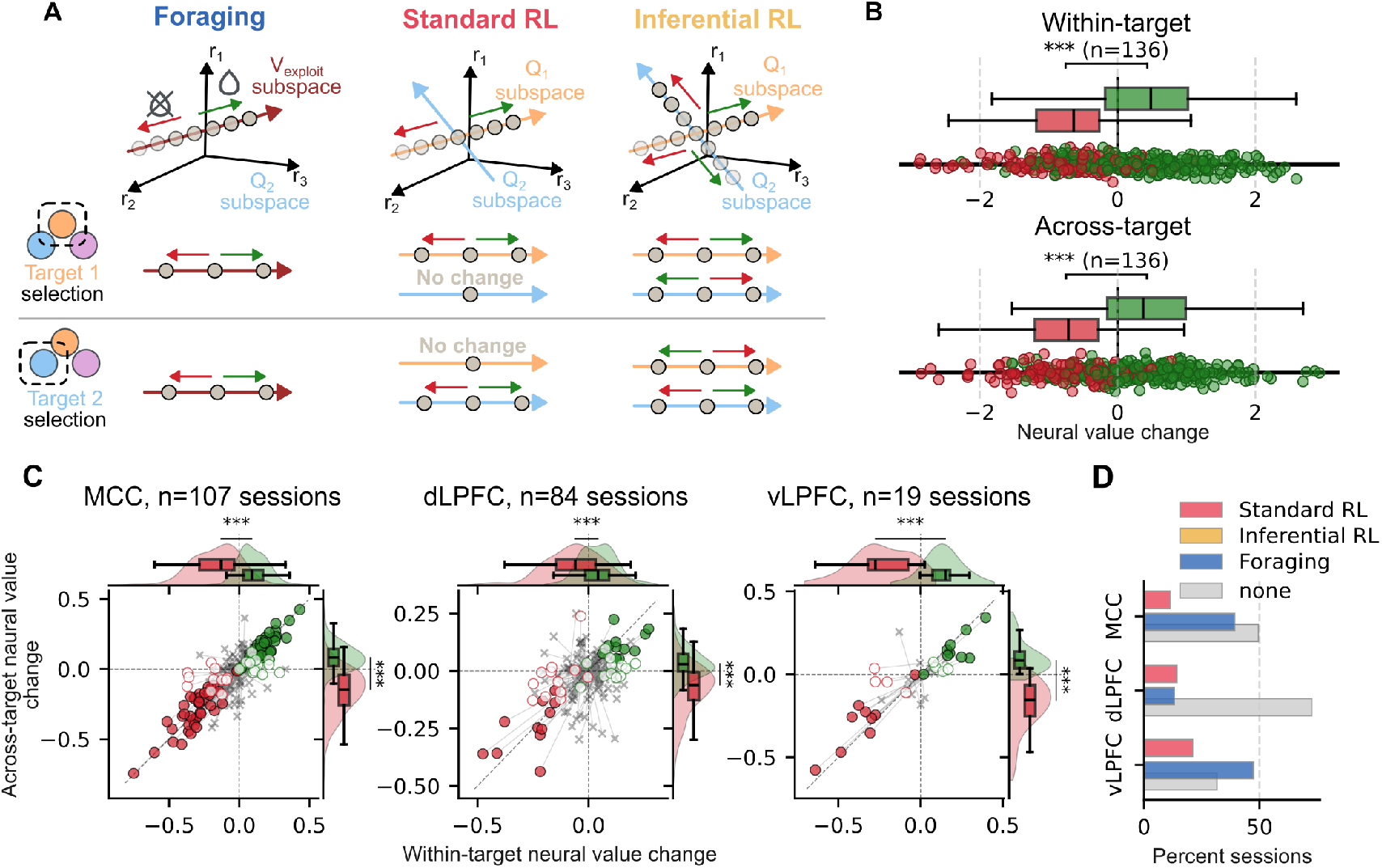
Neural representation of value supports the foraging strategy over RL. **A**, Schematic representation of foraging and RL predictions (columns). Top: firing rate responses (r) of three neurons and DVS of neural value (dark red arrow) for the first target (dashed line on the middle panel). Positive and negative outcomes for that target induce systematic changes in the DVS (bottom panels). Different models predict different outcome-dependent changes in the DVS when the selected target differs from the one the DVS was established for (across-target changes). **B**, Within-target (top) and across-target (bottom) changes in the DVS in an example session. Dots represent trial-by-trial changes in successive trials. **C**, Session-by-session average (each session is represented by a pair of dots for positive and negative outcomes) of within-target and across-target change in the DVS. Colors show outcome valence, symbol is determined by within-target decodability of feedback (cross indicates non-significant decoding performance). Filling of circles indicates if the across-target decoding of outcome is significant. Circles in the SW and NE quadrants are predicted by foraging. **D**, Fraction of sessions providing support for each hypothesis. Note that sessions supporting standard RL might still be compatible with foraging, but not the opposite.

We then examined changes in this neural value signal from one trial to the next under two conditions: in the within-target condition, trials with a given chosen target are projected into the DVS obtained from trials with the same chosen target; in the across-target condition, trials with a different chosen target are projected to the DVS of alternative targets. The foraging model predicts a value representation which is independent of the selected target. Therefore, both within- and across-target conditions should show positive changes in neural value following positive outcomes, and negative changes following negative outcomes. In contrast, reinforcement learning models predict target-dependent value representations: standard RL predicts a positive modulation of value only within-target, with no change across-target, while inferential RL predicts an inverse modulation across-target (**Fig. 5A**).

Analysis of individual sessions revealed reliable modulation of value by feedback, both within- and across-target conditions (see an example session in **Fig. 5B**). The significance of the difference of the change in neural value between positive and negative outcomes was assessed (two-tailed Mann-Whitney U-test, p < 0.05). The test also allowed us to determine the direction of modulation. For each session, we plotted the mean change in neural value following positive and negative feedback in the within-target condition against the across-target condition (**Fig. 5C**). At the population level, all recorded areas showed significant positive modulation in both the within- and across-target conditions (Mann-Whitney U-test, *p <* 0.001). In the MCC and vLPFC, the majority of sessions exhibited target-independent value representation (filled circles in **Fig. 5C**), consistent with the foraging hypothesis, while a smaller proportion of sessions showed target-dependent value coding (empty circles), supporting standard RL mechanisms (MCC: 39.3% foraging & 11.2% RL; vLPFC: 47.4% Foraging & 21.1% RL) (**Fig. 5D**). Note, however, that sessions supporting the standard RL account exhibited weaker value modulation in the within-target condition, suggesting that these effects may lie near the threshold of statistical significance. Importantly, no sessions in any of these three regions supported the inferential RL hypothesis. The dLPFC exhibited a smaller proportion of sessions with significant value modulation overall, with a roughly equal proportion supporting Foraging and standard RL models (13.1% Foraging, 14.3% RL). Results were consistent across monkeys (see **Supplementary Fig. S6** for per-monkey results).

### Temporal evolution of outcome integration

While the previous analysis established precise trial-by-trial updating of the value signal, a key question remains about how this updating unfolds over time. To address this, we performed Principal component analysis (PCA) on the composite session data and, for each principal component (PC), measured activity during the task-engagement window (specifically, during the 2-second period between lever touch and feedback). We then quantified how much of the variance in each PC’s activity was explained by the model-derived value, and how much of the variance in its trial-to-trial change was explained by the received outcome (using one-way ANOVA, see Methods; Fig. 6A). In both monkeys, within the MCC, we identified a single PC most strongly related to both value coding and its updating (**Fig. 6A**, circled markers on left panel). PC components involved both in value and update coding were absent in the dLPFC and vLPFC in either monkey (**Fig. 6A**). These relevant PCs in MCC exhibited stable representation of the value signal, starting shortly after the feedback delivery and persisting through the inter-trial period until the next feedback (**Fig. 6B**). Importantly, value representations updated systematically after the feedback onset: for low initial values, positive outcomes shifted the activity upwards (red plain line), whereas negative outcomes produced minimal change (red dashed line)(**Fig. 6B**). The opposite pattern was observed for high initial values, with large downwards updates following negative outcomes but minimal change after positive outcomes. Intermediate values were influenced by both outcomes.

**Figure 6.**
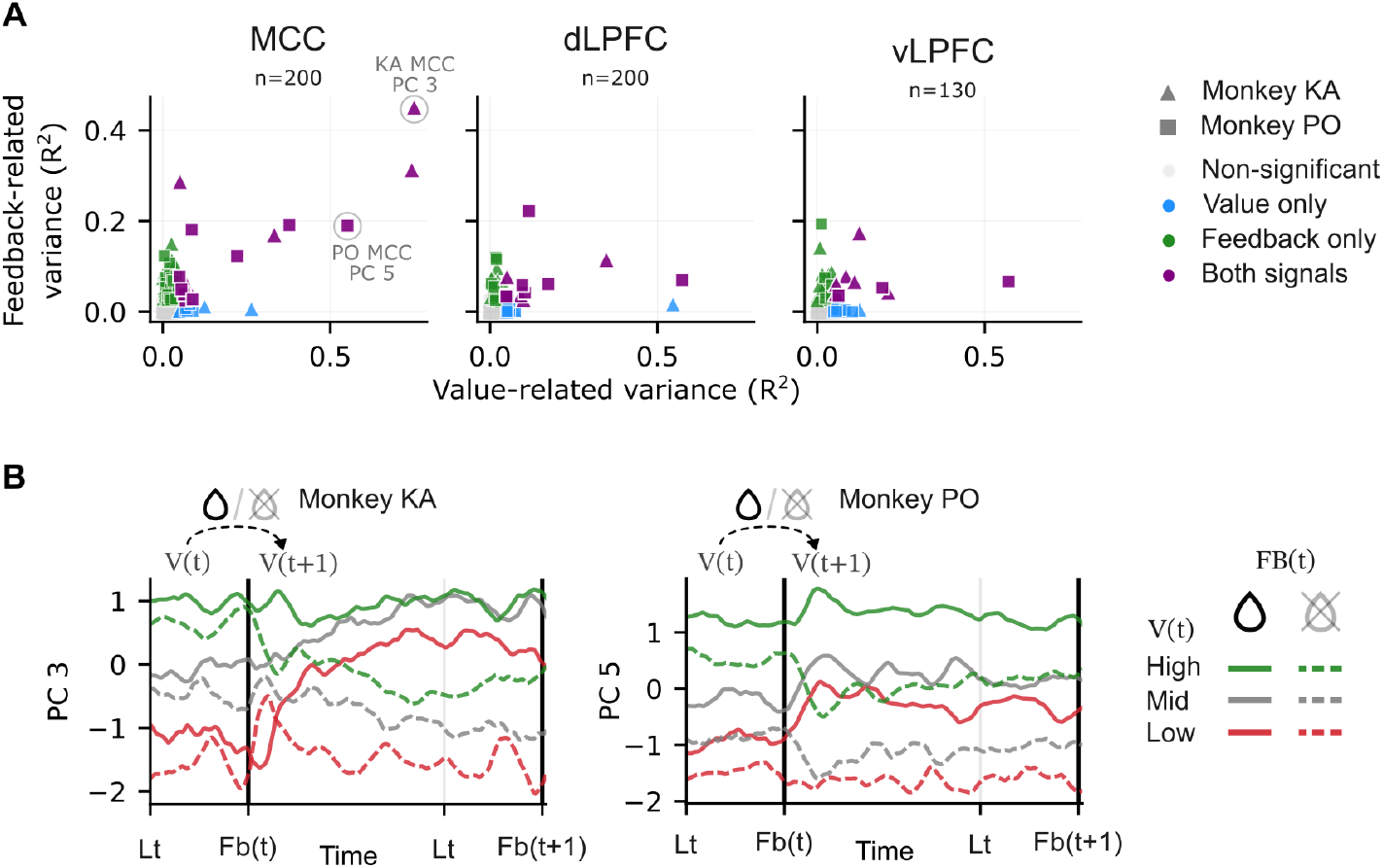
Dynamics of neural value updating in MCC. **A**, PCA was applied to the composite session data. For each principal component (markers: triangle for monkey KA, rectangle for monkey PO), we quantified how much of the variance in activity was explained by model-derived value, and how much of the trial-to-trial variance in activity was explained by outcomes. Significance was assessed per PC and analysis (one-way ANOVA, at p<0.05, blue: variance in activity explained by value, green: variance in trial to trial change in activity explained by feedback, purple: both significant, grey: neither significant). **B**, The linear subspace most correlated with neural value in a PCA decomposition of population responses and population trajectories in this subspace in MCC for both experimental animals. Population trajectories are stratified according to the neural value in a given trial and according to the outcome of that trial. Population trajectories are followed throughout the next trial until feedback is delivered.

## Discussion

We recorded neuron population activity from multiple prefrontal regions of macaque monkeys while they performed a three-armed bandit task, the dominant testbed for reinforcement learning. Through quantitative evaluation of behavior, we could precisely track variables critical for computations in neural activity. By establishing the arithmetic of value guided choice in neural populations, we could contrast predictions of reinforcement learning and foraging. Our recordings showed strong alignment with the foraging framework but did not support a reinforcement learning account. The foraging account supports a precise but simple population code that meticulously tracks the value of the actual choice, comparing it to a threshold to assess the goodness of the choice and dynamically updating the value as new evidence becomes available. Moreover, our analyses revealed select but clear functional dissociations. Both MCC and vLPFC showed signatures of value representations, but only in the MCC could we find evidence for a clean subspace that, in line with foraging, reflects the maintenance and trial-by-trial dynamical update of the value. Our analyses demonstrated that the identified neural code for value representation was optimal as the value estimates obtained from behavior precisely predicted the across-trial dynamics of the value estimated from neural data. In summary, our results demonstrate that the MCC precisely performs a resource-efficient computation to maximize reward.

### Prefrontal functional and dynamical dissociations

The decoding of key variables was applied on neural activity recorded from extensive regions of the MCC, dLPFC and ventral region of LPFC (vLPFC). Recordings in MCC likely included the rostral hand and the face/outcome cingulate areas as described from previous meta-analyses (Procyk et al., 2016). The most rostral sites potentially included the transition between MCC and ACC as suggested by recent anatomo-functional investigations (Ducret et al., 2024). These cingulate regions were associated, both in human and non-human primates, with integrating outcome history or representing task values necessary for regulating behavioral flexibility (Amiez et al., 2006; Kennerley et al., 2006; Seo and Lee, 2007; Kolling et al., 2016; Procyk et al., 2021). Value integration in MCC seemed to go hand in hand with a contribution in switching, e.g. from default states to exploration, foraging, or information seeking (Quilodran et al., 2008; Monosov, 2017; Kolling et al., 2012, 2018; Kolling and O’Reilly, 2018).

Our study showed clear functional specificity between frontal subregions. Both MCC and vLPFC revealed trial-to-trial neural changes reflecting value updating. vLPFC recordings were within the ventral border of the sulcus principalis, in area 46v or 46vc (Borra et al., 2011, 2019). Although limited in number, these sites displayed functional properties very similar to those found in MCC, and in fact both regions are anatomically connected (Gerbella et al., 2013). Our data thus suggest an integrated role for the MCC-vLPFC (46v) network in monitoring performance and switching, which might be coherent with recent propositions regarding MCC interactions with more ventral regions of vLPFC (Monosov and Rushworth, 2022). MCC and vLPFC neural encoding clearly differed from dLPFC. Note that recordings covered a large rostro-caudal extent of the dLPFC territory that has been targeted across past neurophysiological studies (see fig. S11 in Amiez et al. (2023)). This corresponds to the caudal, middle, and a part of rostral 46d as described by Borra et al. (Borra et al., 2019). Here we did not segregate between subregions of the dLPFC although many studies observed fine functional dissociations for shifting between abstract sequences, neural timescales, and feature dimensions in working memory (Rodriguez et al., 2024; Riley et al., 2017; Trepka et al., 2024). We show here that dLPFC neural dynamics mostly track target selection and identity (spatial position) during the 3-arm bandit task, but with little if any trace of value coding and updating.

The regional specificities we described are somewhat aligned with previously observed dissociations (Kennerley and Wallis, 2009; Hunt et al., 2018; Seo and Lee, 2007), but they also substantially refine known neurophysiological and mechanistic dissociations, specifically regarding value integration. Several past studies have shown that neural intrinsic timescales vary between medial and lateral subdivisions of the frontal cortex. The variations contribute to the computational specificity of cortical subdivisions, notably by changing dynamical properties and metastability that might help information integration across multiple timescales (Fontanier et al., 2022; Cavanagh et al., 2018; Bernacchia et al., 2011). In fact, MCC activity was consistently shown to express longer timescales (Fontanier et al., 2022; Murray et al., 2014).

The particular relevance of MCC for integrative functions in the context of reinforcement learning, for instance, was suggested by lesion, computational, and neurophysiological studies (Khamassi et al., 2015; Cai and Padoa-Schioppa, 2012; Kennerley et al., 2006). Also, in contrast to LPFC and OFC (orbitofrontal cortex), MCC neural activity specifically correlates with belief updating during sequential information sampling (Hunt et al., 2018). Here we show that the integration of outcomes in a value used for decisions to switch is encoded in multiple areas of the frontal cortex, specifically in MCC and vLPFC. However, the updating of value according to a delta-rule was selectively observed in MCC. Moreover, the population decoding revealed that neural dynamics were better explained by a straightforward foraging model compared to action-outcome learning models.

Our analyses allow us to present a mechanistic account of foraging-like decision making in line with previous findings on MCC function. The model relies on two distinct computations: (1) valuation: inferring value from past experience via standard error-based learning; and (2) choice: comparing the running value to a leave threshold to decide whether to persist or switch. Essentially, the model accumulates evidence only for the currently pursued option, and initiates switching when this value falls below a leave threshold. Modulating the learning rate and the threshold allows the model to adapt to the environment’s reward structure. This connects to MCC single unit activity shown rising to a threshold and correlating with patch leaving decisions during foraging-like tasks (Hayden et al., 2011). White and colleagues suggested that MCC computations might act through an MCC-striatal pathway to motivate actions to explore or seek information (White et al., 2019). More recently, Gandaux et al. showed that the MCC-to-LPFC pathway causally contributes to motivational control of exploratory behaviors (Gandaux et al., 2025). These observations align with theoretical propositions that monitoring and evaluative functions implemented in MCC specify, through larger scale networks, which control to apply in the current situation (Shenhav et al., 2013; Domenech and Koechlin, 2015). Still, little is known about the exact computational and neural mechanisms leading to value integration in MCC and how it could inform or trigger switching decisions. Our study provides an in-depth description of MCC computations for switching decisions and a coherent mechanistic account that unifies MCC functions that were often described separately: temporal evidence integration and control of switching behavior.

### Foraging as model-based control

In our study, we contrasted reinforcement learning (RL) and foraging as potential computational strategies underlying the neural population dynamics. The investigated strategy is a specific form of RL: model-free reinforcement learning (Sutton, 2018; Drummond and Niv, 2020). The theory of RL also distinguishes model-based algorithms; in neuroscience, a major theme is to distinguish model-based and model-free strategies both in animal and human behavior (Collins and Cockburn, 2020; Daw et al., 2005; Frömer et al., 2021; Kahn and Daw, 2025; Keramati et al., 2011; Kool et al., 2017; Piray and Daw, 2021; Eckstein and Collins, 2020; Daw, 2018; Drummond and Niv, 2020; Daw et al., 2011). While model-free RL assigns values to state-action pairs, a model-based algorithm also learns potential transitions among states and evaluates alternative policies over the potential transitions. A model-based RL algorithm is more flexible as it enables efficient reuse of learned information to generalize to unseen situations (Drummond and Niv, 2020; Gershman and Daw, 2017). Interestingly, a foraging strategy can be cast as a model-based hierarchical reinforcement learning (HRL) (Alejandro and Holroyd, 2024) by treating harvest grounds (patches) as options and having a high-level controller plan for leaving the grounds or staying. Interpreted as a HRL, in our task, high-level switch/stay decisions gate low-level action selection. In model-based HRL strategies, a foraging strategy can be interpreted as a naturalistic prior that the animal recruits when faced with novel environments. Recent evidence supports that humans implement HRL (Eckstein and Collins, 2020). In the context of interpreting foraging as HRL, recent accounts explicitly situate the cingulate cortex (in particular the MCC) as a controller within a model-based HRL architecture (Alejandro and Holroyd, 2024). In summary, our results are compatible with a model-based reinforcement learning setting but are at odds with a basic model-free interpretation. Consequently, even when model-free algorithms can approximate near-optimal performance, animals likely deploy ecologically tuned, model-based priors when facing simple economic tasks, suggesting that simple model-free accounts – common in lab tasks – can miss the policy the brain actually uses (Daw et al., 2011; Constantino and Daw, 2015; Alejandro and Holroyd, 2024; Zid et al., 2025).

### Generalized computations in a foraging context

A hallmark of decision making is that the conditions under which the decisions are made introduce uncertainty at multiple levels, not only about the value of the choice, but also about the value of alternatives, or the properties of the environment. Probabilistic computations have been explored in a wide array of contexts, including perception (Orbán et al., 2016) and planning (Ujfalussy and Orbán, 2022). More recently, a probabilistic version of RL, distributional RL (Bellemare et al., 2017) has been proposed for decision making under uncertainty, and neural signatures of such probabilistic computations have been identified in midbrain dopamine populations (Dabney et al., 2020) and in prefrontal cortex, including regions overlapping the MCC (Muller et al., 2024). We have not explored the consequences of uncertainty in the present work, but integration of the foraging account presented here and the probabilistic components of distributional RL could provide additional clues into the precise computations taking place in the MCC.

## Resource availability

### Lead contact

Requests for further information and resources should be directed to and will be fulfilled by the lead contact, Emmanuel Procyk.

### Materials availability

This study did not generate new unique reagents.

### Data and code availability

Data reported in this paper will be shared by the lead contact upon request. All code required to reanalyze the data reported in this work will be available upon publication of the paper.

## Acknowledgements

We would like to thank Nils Kolling for his valuable feedback on our manuscript.

## Author contributions

Conceptualization, Z.U., C.G., A.S., G.O., M.d.V., E.P.; Data curation (preprocessing), C.G., Z.U.; Formal analysis (main results), Z.U., A.S.; Funding acquisition, G.O., M.d.V., E.P.; Investigation (recordings), C.G.; Methodology (task design), C.G., E.P.; Methodology (main analyses): Z.U., A.S., G.O., M.d.V., E.P.; Supervision (main analyses), G.O., M.d.V., E.P.; Supervision (task design/recordings/preprocessing), E.P.; Visualization, Z.U.; Writing – original draft, Z.U., G.O., E.P.; Writing – review & editing, G.O., M.d.V., E.P.

## Methods

### Experimental setup

Ethical permission was provided by “Comité d’Éthique Lyonnais pour les Neurosciences Expérimentales,” CELYNE, C2EA#42, under reference: APAFIS#14704_2018041318163080. Monkey housing and care was done in accordance with the European Community Council Directive (2010) and the Weatherall report. Laboratory authorization was provided by the “Préfet de la Région Rhône-Alpes” and the “Directeur départemental de la protection des populations” under Permit Number: #B-690290402. Data were collected from two rhesus monkeys: one male (monkey PO, 11 years old) and one female (monkey KA, 12 years old).

### Behavioral task

Monkeys performed a three-armed bandit task in which they identified, through trial and error, which target among three options offered the highest probability of reward (**Fig. 1A-C**), a 1:1 mixture of water and apple juice. In this three-armed bandit task, one of the three targets yielded a reward 70% of the time, while the other two targets each provided rewards on 25% of presses. The highest rewarded target changed from block to block. The sequence of targets was randomly determined at the beginning of each session. Each block consisted of 40 trials on average, ranging from 35 to 45, sampled randomly. Reward probabilities transitioned gradually during the last five trials of a block.

Monkeys had to follow a specific sequence of actions: first, the lever had to be pressed (lever touch) and held for 1000 ms to validate it (lever validation). Second, one of the three displayed targets had to be pressed (target touch) and held for 450 ms to validate a choice (target validation). The outcome, i.e. either juice (rewarded) or no juice (unrewarded), was obtained 500 ms after this sequence (feedback). Each trial concluded with an inter-trial interval (ITI) of 2 to 3 seconds, set randomly. Monkeys had to release their hand from the screen during this interval to initiate a new trial.

Monkeys interacted with a touch screen in a sound-attenuated box. To suppress environmental auditory noise, a continuous red noise was played throughout the recording period. The visual layout on the touch screen featured the three targets arranged around the central lever, and remained displayed throughout the task, mimicking the stability of real-world objects. Shadows and depth cues were added to the stimuli to simulate the appearance of natural three-dimensional buttons (**Fig. S2A**). When a monkey touched either a target or the lever, the visual elements dynamically changed to resemble the pressing of a physical button. Unless already validated, both the lever and the target returned to their initial (untouched) position when monkeys released them. Successful validation of both the lever and the targets was indicated by a sound signal. If validated, a yellow bar appeared around the lever (**Fig. S2A**). At the end of the ITI, a new trial initiated only if the monkey released the screen. This was accompanied by a sound signal (distinct from the validation signal), indicating that the lever returned to its original unpressed state and was once again active.

Visual display, control of the task schedule, and behavioral monitoring was managed using the EventIDE software (OkazoLab, Netherlands). Monkeys performed the task using their left hand.

### Electrophysiological recordings and spike sorting

#### Surgical procedures

Head post and recording chamber were implanted in separate surgeries. The titanium recording chamber (20mm x 20mm, Rogue Research Inc., Canada) was fit to each individual cranium thanks to T1 MRI 3D reconstructions, and was implanted using neuronavigation (Brainsight, Rogue Research Inc.). The chamber was implanted over the hemisphere contralateral to the arm used. The ideal position of the chamber was calculated taking into consideration both previous recordings, meta-analysis and the particular morphology of each brain (Procyk et al., 2016). Per-surgery medication to avoid pain and inflammation were used. Analgesics and antibiotics were administered throughout surgical periods. The chamber was kept sterile with regular cleaning and sealed with sterile caps.

#### Recordings

Recordings were performed simultaneously in the lateral prefrontal cortex (LPFC) and the midcingulate cortex (MCC). Specific location of recording sites is shown in **Fig 3A** for monkey KA and **Fig. S2B** for monkey PO. MCC recordings were located in the dorsal bank of the cingulate sulcus. In each session, before recording began, two V-probes (V-probes; Plexon Inc., Texas, USA), each with 16 contacts spaced 200 µm apart, were slowly (3 µm/s) and perpendicularly inserted into the targeted area. During each recording, one probe was driven into the LPFC and the other into the MCC. Electrodes were advanced orthogonally into the LPFC and MCC at a controlled rate using a Flex microdrive (AlphaOmega eng., Israel). This procedure lasted approximately one hour and recording started once the neural signals were stable in time. During a recording session, the monkeys were placed in a specific chair designed for behavioral and neural recording. To ensure stability, the monkey heads were held in place via a preimplanted head post.

Electrical referencing was achieved via connections to both the head chamber and the electrode tip, while grounding was performed externally. A 96-channel Cereplex Direct acquisition system (Blackrock Microsystems, USA) was used to record neural signals, capturing broadband activity between 0.5 Hz and 9 kHz at a sampling rate of 30 kHz. This setup enabled synchronized acquisition of neural, behavioral, and eye tracking data within a unified temporal framework.

The collected data was pre-processed using the Blackrock and FieldTrip toolboxes for Matlab, and the KiloSort package with Python back-end. Spikes were extracted from the recordings using KiloSort v2 (Pachitariu et al., 2016). The output of KiloSort was then manually curated with the help of *Phy*, a Python library allowing for visual correction and manipulation of clustered data. Spike times were down-sampled to 1000 Hz by rounding each spike time to the nearest 0.001 s.

### Behavioral analysis

We analyzed behavioral data collected over 62 and 55 sessions, consisting of 9.1*±*1.7 and 6.6*±*2.5 and 6.6*±*2.5 blocks on average, of monkey KA and PO respectively. In total, our analysis included 23,952 trials for monkey KA and 15,267 for PO. Interrupted trials, during which monkeys stopped interacting with the task, were excluded from the data analysis (KA: *n* = 15, 0.06%; PO: *n* = 29, 0.19%).

We computed two behavioral metrics to visualize within-block dynamics: (i) the percentage of trials where the animals selected the target with high payoff, and (ii) the probability of switching to the alternative option (**Fig. 1D-E**). In each block, we extracted the series of binary variables of interest (e.g. optimal target or not; switch or stay). To align series across blocks with different lengths, each were interpolated to a length of 40 trials. For each monkey separately, we extracted the proportion (mean) and standard error of the mean (SEM) of the variables of interest across blocks for each aligned trial position.

We used a logistic regression model to predict whether the animal would switch to an alternative option (as opposed to staying with the previously selected option, based on the history of the last 10 outcomes (**Fig. 1F**). The model thus took the form of

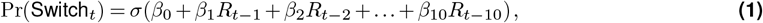

where Pr(Switch_*t*_) is the probability to switch to an alternative option at trial *t, β* values are regression coefficients, the *R*_*t−i*_ denotes the outcome of the *t ™ i*-th trial (coded as 0 for non-rewarded and 1 for rewarded) and *σ*(·) is the logistic function. To account for the imbalance between switch and stay trials, the model was fit using class-balanced weights. We implemented this model using the scikit-learn library (Pedregosa et al., 2011). For each monkey, coefficients were estimated from the full dataset, and statistical significance was computed using nonparametric permutation test: switch/stay labels were randomly shuffled 1000 times, the model was refit and weights were assessed, then empirical two-sided *p*-values were computed as the proportion of permuted coefficients exceeding the true observed coefficient. Predictive accuracy of the model was evaluated with 10-fold cross validation.

### Decision making models

#### Win-Stay-Lose-Switch model

As a baseline, we implemented a win-stay-lose-switch (WSLS) model, that determines whether to switch or stay based on the last three outcomes. This model implements a simple decision rule: it repeats the previous action if at least one of the last three outcomes was rewarded and switches otherwise (i.e., if all three of the most recent outcomes were unrewarded). The WSLS model provides efficient performance under minimal computational cost.

Action selection was implemented as a noisy function of the switch-or-stay strategy: at each trial, the stay (switch) strategy was followed with a probability of 1 *− ϵ*, while the opposite switch (stay) strategy was executed with a probability of *ϵ*. Thus the probability of choosing each action *a*_*k*_ *∈ {a*_1_, *a*_2_, *a*_3_*}* in trial *t* can be formally expressed as

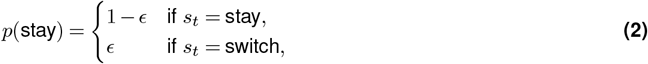

where *s*_*t*_ ∈ {Stay, Switch} is the corresponding strategy: Stay if at least one of the three most recent trials was rewarded, and Switch otherwise.

Accordingly, the probability of selecting each action *a*_*k*_ *∈ {a*_1_, *a*_2_, *a*_3_*}* at trial *t* is

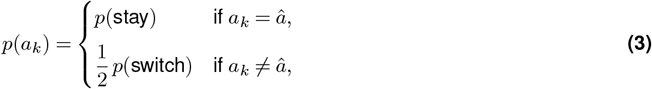

where *p*_Switch_ = 1 *− p*_Stay_ and *â* represent the previously selected action. The WSLS model contains a single free parameter, which controls the overall randomness of the decision: *θ*_WSLS_ = *{ϵ}*.

#### Reinforcement learning models

The standard solution for a three-armed bandit task is a Q-learning model (Sutton, 2018), referred to as the *standard RL* model in this work, which assigns one value to each action (arm) and uses these action-values to select an action. The standard RL model updates the value of the chosen action *a*_*k*_ in response to the reward *R* according to the delta learning rule but does not update the values of the unchosen actions:

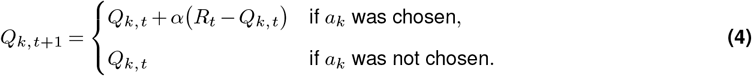

where *α* is the learning rate, and *R*_*t*_ is the reward (unrewarded outcome = 0, rewarded outcome = 1). For action selection, Q-values are transformed to action probabilities using the softmax function, with an additional *stickiness parameter κ*, controlling the probability to repeat the previously selected option (*κ >* 0 results in repeating the last action more often). The probability of choosing each action *a*_*k*_ *∈ {a*_1_, *a*_2_, *a*_3_*}* in trial *t* is given as

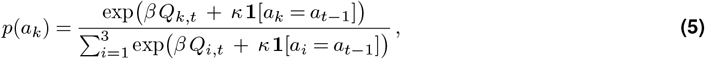

where *β* is the inverse temperature parameter, which controls the degree of randomness in the decision, and **1**[·] is the indicator function (1 if the condition is true and 0 otherwise). The Q-learning model has three free parameters: *θ*Standard RL = {*α, β, κ*}.

We used a modified version of the Q-learning algorithm, which we refer to as the *inferential RL* model (**Fig. 1H**), which accounts for the anticorrelation structure of the targets – i.e., that there is always one and only one target with high probability and consequently the two others have low probabilities. This means that information about one action can be used to update the values of the other actions. The model updates the values of the unchosen arms with inverse reward. Thus the value of each action *a*_*k*_ *∈ {a*_1_, *a*_2_, *a*_3_*}* in trial *t* updates as

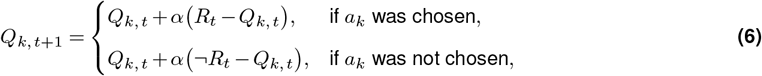

where ¬*R* is the inverse reward (unrewarded outcome = 1, rewarded = 0). Similar to the standard RL, this model also uses the softmax choice rule with stickiness (Equation 5) for action selection. The inferential RL model has thus two free parameters: *θ*_Inferential RL_ = *{α, β, κ}*.

#### Foraging model

The foraging model (**Fig. 1G**) integrates reward history into a single quantity representing the value of continuing to exploit the currently chosen action, *V*_exploit_. This value is compared to a *switch threshold* (*τ*) to decide whether to further exploit the currently selected action or switch to an alternative. *τ* reflects the global reward statistics of the environment (i.e., the long-term average reward rate), which predicts the value of random exploration. We assume *τ* is learned during extended training, thus we treat it as a fixed threshold.

The value *V*_*exploit*_ updates trial-by-trial based on the delta learning rule, thus at trial *t* + 1 it is calculated as

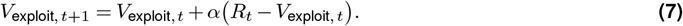

Following switching, the value *V*_*exploit*_ is reset to the baseline *τ*, consistent with a patch-leaving interpretation (Charnov, 1976). *V*_*exploit*_ is then compared against the value of exploration, *τ*, using the logistic function to determine the probability of exploiting or switching:

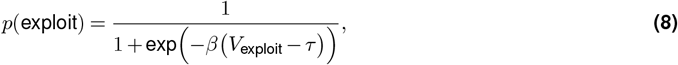

where *β* controls the level of noise in the decision. In short, the difference between the value of exploitation and the switching threshold, *V*_*exploit*_ *−τ*, determines switching probability.

Accordingly, the probability of selecting each action *a*_*k*_ *∈ {a*_1_, *a*_2_, *a*_3_*}* at trial *t* is

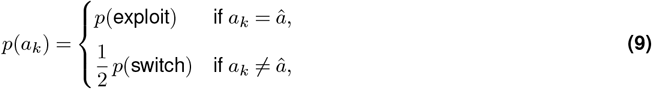

where *p*(switch) = 1 *−p*(exploit) and *â* represent the previously selected action. The foraging model has three free parameters, *θ*_foraging_ = *{α, β, τ}*.

#### Hidden Markov Model

To provide an optimal account of choice in our task, we implemented a Hidden Markov Model (HMM). The model casts the bandit problem as a latent-state model with three states, where in each state one of the targets has a high payoff probability and the other two have low payoff probabilities. The HMM was given exact knowledge of the task’s generative structure: it maintains an emission matrix for each state (*p* = 0.7 for the high-payoff arm, *p* = 0.25 for the rest) and a transition matrix that represents volatility (i.e., a 1*/*40 chance that the state switches to one of the two alternatives). The model infers the true state of the environment based on its prior belief of the states and the current outcome, using Bayesian inference. It accounts for the volatility of the environment by projecting the posteriors forward through the transition probabilities. Posterior beliefs together with the emission matrix provide predictions of the expected outcomes of each choice.

On each trial, first, the posterior probability of states is updated based on the observed reward *R*_*t*_ and the chosen action *a*_*t*_, using the emission probabilities:

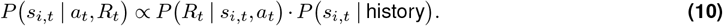

where *s*_*i,t*_ denotes the latent state *i* ∈ {1, 2, 3} at trial *t, P* (*R*_*t*_ |*s*_*t*_, *a*_*t*_) is given by the emission matrix, and *P* (*s*_*t*_ | history) is the prior.

Second, the posterior is updated with the state transition matrix, which corrects the posterior for the volatility of the environment.

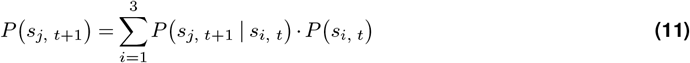

Action selection is based on the posterior distribution over states: the probability of selecting *a*_*k*_ ∈ *a*_1_, *a*_2_, *a*_3_ is given by a softmax transformation over the marginal reward probabilities, weighted by the posterior belief, which in our case is equivalent to a softmax over the posterior (as the emission matrix is symmetric), up to a rescaling of *β* inverse temperature parameter:

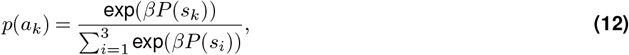

where *s*_*k*_ represents the state corresponding to *a*_*k*_ (in which *a*_*k*_ has a high payoff). The HMM has one free parameter: *θ*_*HMM*_ = *{β}*.

#### Optimizing for performance

To validate that the models were able to reproduce monkeys’ performance in the task, we fit their parameters to maximize the average reward during simulations (**Fig. 2A**). The performance-optimal parameter set, 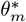 for each model *m*, was then given as

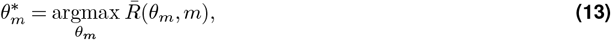

where 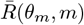 is the mean reward per trial obtained by model *m* with parameters *θ*_*m*_ during a simulated run of 100,000 trials. We used a Gaussian Process–based Bayesian optimization procedure to find the performance-optimal parameters for each model, which minimized the negative average reward as the objective function (implemented with <monospace>scikit-optimize</monospace> (Head et al., 2020).

#### Model fitting

Models were fit using the Maximum Likelihood Estimation (MLE) method to find parameters, *θ*_*m*_, for each model, *m*, that best described the behavioral data. Formally, we aimed to obtain optimal parameters, 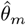, to maximize the log-likelihood, as

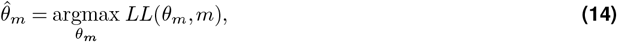

where the log-likelihood function was given as

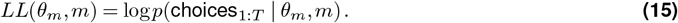

To find the maximum likelihood, we minimized the negative log-likelihood using a Gaussian Process–based Bayesian optimization procedure (Head et al., 2020).

We quantified how model-derived values predicted choices using the receiver operating characteristic (ROC) analysis and area under the ROC curve (AUC) (**Fig. 1J-M**). Value estimates were computed for each trial using the best-fitting parameters (see Model Fitting). The foraging model produces a single value (*V*_exploit_) that predicts staying tendency. Thus, a simple binary classification of choice (switch or stay) against the value was evaluated with the ROC-AUC. The inferential RL model represents one action-value signal per target ({*Q*_1_, *Q*_2_, *Q*_3_}) and predicts directly the selected target. Thus we evaluated multiclass prediction using one-vs-rest approach. We computed binary ROC curves for each value separately, and we reported macro-averaged ROC-AUC across the three one-vs-rest ROCs (implemented with the scikit-learn library (Pedregosa et al., 2011)).

We compared models to determine which was most likely to have generated the data. For each model *m* and its corresponding best parameters 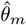, we computed the likelihood-per-trial (LPT), defined as the average log-likelihood per trial (**Fig. 2B**). Given that models have at most three free parameters and behavior data contains at least 10,000 trials, the effect of correcting for complexity (e.g., Bayes Information Criterion) would be negligible – we instead implemented the more intuitive LPT measure. Model fitting was estimated using 3-fold cross-validation.

### Analysis of neuronal responses

To characterize neural encoding of task variables at the single unit level, we analyzed trial-by-trial variations in spike counts using generalized linear models (GLM). We used spike count data for the analysis, relying on 200-ms long windows, sliding in 50-ms intervals. This smoothing allowed us to reduce trial-to-trial variability in firing rates, while preserving sufficient temporal resolution to track the dynamics of neural encoding. Time labels mark the middle of this moving window, e.g. a data point at *t* = 500 ms would include data from a window between 400 and 600 ms.

We analyzed neural activity in a 6-s window centered on feedback.

Neural responses were predicted using standard Poisson GLM. The model predicted spike counts based on the Reward (binary: 0/1) and the model-based value of the foraging model (continuous: 0-1)

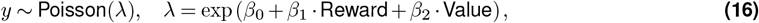

where *y* represents the spike count in a given time bin, *λ* is its expected value, and the coefficients *β* were fit without regularization using the statsmodels package (Seabold and Perktold, 2010).

To quantify the contribution of individual regressors to the performance of the full model, we computed the Coefficient of Partial Determination (CPD) for each regressor. CPD measures the percentage of increase in explained variance when the relevant coefficient is included:

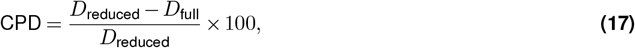

where *D* is the mean Poisson deviance, computed for the model with all regressors (*D*_full_) as in Equation 16, and for the reduced model where the relevant regressor was omitted (*D*_reduced_). The mean Poisson deviance is calculated by averaging across trials the Poisson deviance between observed spike counts *y*_*t*_ and predicted rates *ŷ*_*t*_:

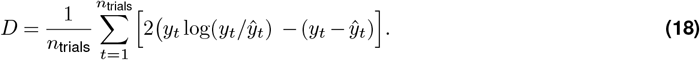

The CPD was averaged over neurons at each time bin to give population-wise statistics.

We used permutation testing to evaluate the statistical significance of the resulting CPD values. The dependent variable was randomly shuffled and the CPD was computed 500 times to approximate its null distribution. p-values were then calculated as the proportion of permuted CPDs greater or equal to the CPD of the observed data. The significance criteria was set to *p* ≤ 0.01. Single units were classified as coding for a variable if they showed a significant CPD (*p <* 0.05) in at least four consecutive time bins. Feedback coding was tested in a 1000-ms time window after feedback delivery; value coding was tested in a 1000-ms window prior to feedback delivery.

### Firing rate generation

For the population-level analyses, spike times were converted to firing rates by smoothing the raster of each cell with a Gaussian kernel (*σ* = 50*ms*). Due to the instability of the recording electrodes, there were neurons that we were unable to track for the entire duration of the session. For population-level analyses, we required neurons to exhibit stable activity during the session. Activity was considered stable if both the first and last spike occurred within 20 s of the recording start and end, respectively. As a consequence, analyses of single units include more neurons compared to population-level analyses.

### Time normalization

In our paradigm, the length of each trial differed as the length of certain epochs (between trial start to lever touch; and lever validation to target selection) depended on the response time of the animals. Inter-trial interval, that is the epoch after the feedback delivery also varied (see Methods, **Behavioral task**). In order to be able to describe the evolution of the neural activity throughout the whole trial, we normalized time in epochs to obtain uniform trial length across trials. Specifically, we resampled firing rates in each epoch to obtain *n* time bins (st → Lt: *n* = 100, Lt → Lv: *n* = 100, Lv → Tt: *n* = 50, Tt → Tv: *n* = 50, Tv → Fb: *n* = 50, Fb → st: *n* = 400), representing 1 s, 1 s, .5 s, .5 s, .5 s, and 4 s, respectively, under the assumption of 100 Hz sampling rate.

### Population decoders

Key behavioral variables were decoded from the population activity using linear decoders, by sliding the decoder in 100 ms steps. Prior to analysis, all units with mean firing rate ≤ 1 Hz were excluded. Firing rate vectors for each unit were *z*-scored. We decoded Feedback (binary: 0/1), Target (multi-class: 1/2/3), and Value (continuous: 0–1) (**Fig. 3F**). Logistic regression with L2 regularization was used for Feedback, one-vs-rest logistic regression with L2 regularization for Target, and Ridge regression for Value decoding. Classes were balanced with random under-sampling for Feedback and Target decoding. All decoders were tested using 10-fold cross-validation.

### Building composite sessions

We pooled neural firing rates together into a “composite session” to give an interpretable view of coding dynamics. To do so, we aligned trials across all units (and thus across sessions) within each monkey and brain area, based on feedback history or value history. Thus, two “composite sessions” were compiled: (i) based on the sequence of the last three outcomes, and (ii) based on the value (grouped into four levels) and the outcome. For each neuron, *n* = 20 trials per class were randomly sampled. Neurons with less than *n* trials in any class were excluded. The selected trials were then concatenated to construct a compound session with dimensions *n*_units_ *× n*_trials_ *×* time. In both composite sessions, eight classes were defined (eight different outcome combinations of three-trial horizon; four value levels x two outcomes; (i) and (ii) respectively), resulting in *n*_trials_ = 160 trials. The resulting compound session consisted of (i) *n*_units_ = 1559 for monkey KA (MCC: 805, dLPFC: 549, vLPFC: 205) and *n*_units_ = 1039 for monkey PO (MCC: 623, dLPFC: 366, vLPFC: 50), and (ii) *n*_units_ = 1166 for monkey KA (MCC: 573, dLPFC: 414, vLPFC: 179) and *n*_units_ = 671 for monkey PO (MCC: 396, dLPFC: 245, vLPFC: 30).

### Decision-vector subspace

Datasets (either single sessions one by one or the compound session) were projected to the subspace defined by the decision vector (DVS) of a logistic regression model (see decoders above). Model fitting and projection were performed with 100 ms resolution. We used 10-fold cross-validation for the projection: we defined the DVS on the training trials and projected only the held-out trials; projected trials were then concatenated across folds. Trajectories were then averaged for each condition defined by the last three outcomes, for visualization. The mean activity in the projection between lever touch (Lt) and feedback (Fb) was defined as the *neural value*.

Modulation of the neural value signal was characterized statistically: modulation by the last three outcomes was assessed with a Spearman’s *rho* test (**Fig. 4C**), and its trial-to-trial changes were assessed using a Mann–Whitney *U*-test (**Fig. 4D**). To quantify trial-by-trial neural value updating, we fit a general linear model separately for each session,

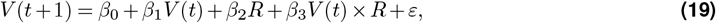

where *V* (*t*) and *V* (*t*+ 1) are neural values from consecutive trials and *R* indicates the outcome (positive or negative). We extracted slopes and intercepts for negative outcomes (given by *β*_0_ and *β*_1_ respectively) and positive outcomes (*β*_0_ + *β*_2_ and *β*_1_ + *β*_3_). We applied three statistical tests: (i) slopes for both outcomes had to be significantly different from zero (two-sided Wald test); (ii) both slopes had to fall between 0 and 1 (one-sided Wald tests 0 and 1); and (iii) the intercept for positive outcomes had to exceed that for negative outcomes (one-sided Wald test). Sessions satisfying all three criteria at *p <* 0.05 were classified as using the delta update rule (**Fig. 4E**).

The modulation of the neural value by outcome history was assessed using the CPD method (see Equation 17), with the full model regressed the neural value, *y*, against the last eight outcomes (binary: 0/1)

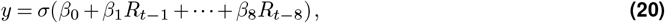

where *σ* is the standard logit function and *β* are the regression coefficients.

We then computed the CPD for each regressor (**Fig. 4F**). Statistical significance was assessed using the same procedure described in section **Analysis of neuronal responses** above.

#### Within- and across-target DVS

We define target-based DVS analysis to characterize the dependence of value representation on the selected target. For this, in each session separately, trials were split into three subsets based on the selected target. In the within-target condition, a target-specific subset was further divided into a training and a test subset: the training subset was used to establish the DVS for that target and the test set was protected into the DVS. As target-wise stratification of the data resulted in limited trading subsets, we chose a leave-one-out cross-validation. In the *across-target* condition, the DVS was fit on one set of trials and the two held-out sets were projected onto it. The *neural value* was then measured in the within- and across-target projections using the method described above. This procedure was repeated for all three targets to obtain one within-target and two across-target neural values per trial. We compared the trial-by-trial change of neural value between the two conditions as a function of the feedback (**Fig. 5B**). For each session, the mean change in neural value was extracted as a function of the outcome for the two conditions (**Fig. 5C**). This analysis reveals how value coding is modulated by action selection.

### Learning-Rate Fitting of Neural Population Activity

We derived trial-by-trial estimates of *V*_exploit_ value of the foraging model with varying *α* learning rates (*α* ranging from 0.05 to 1 with steps of 0.05) to test which best explained variance in neural activity. The same population decoding pipeline was applied session-by-session for each *α*. Resulting *R*^2^ scores were averaged at each time point of the trial to obtain a population-level measure of decoding performance (**Fig. 4G**, left). For visualization, *R*^2^ trajectories were further averaged between lever validation (Lv) and feedback (Fb) per *α* (**Fig. 4G**, right). Only sessions that exhibited significant value coding with *R*^2^ ≥.1 during at least one time bin within the LV–Fb window for any learning-rate parameter were included in the analysis.

### Principal component analysis

A Principal component analysis (PCA) was performed on the compound sessions constructed by aligning trials across four levels of value and the outcome (alignment (ii) in section **Building composite sessions**). For each trial in the composite session, neural activity was taken from a window spanning that trial and the immediately following trial (a time window spanning 15 s in normalized time), in order to capture the dynamics of value updating. Firing rates were *z*-scored, and PCA was fit on the flattened dataset (features = neurons, samples = time points across trials). For each PC, we measured the mean PC signal during the 2-second period between lever touch and feedback onset. We quantified how much variance in this signal was explained by the model-derived value, using a one-way ANOVA with factor ‘value’ (4 levels) applied on the mean PC signal. Feedback-related variance was quantified by first computing the trial-to-trial change in the mean PC signal, and then running a one-way ANOVA with factor ‘feedback’ (2 levels). We report the explained variance as *R*^2^ = SSvalue*/*SStotal and *R*^2^ = SSfeedback*/*SStotal, respectively. Significance levels were assessed using the ANOVA F-test *p*-values (PR(>F) from the model) and is reported in **Fig. 6A**. For visualization, trial-averaged trajectories were computed for each behavioral condition (value levels × outcome) (**Fig. 6B**).

All data analyses were implemented in Python (version 3.10.13).

## Supplementary Information

**Figure S1.**
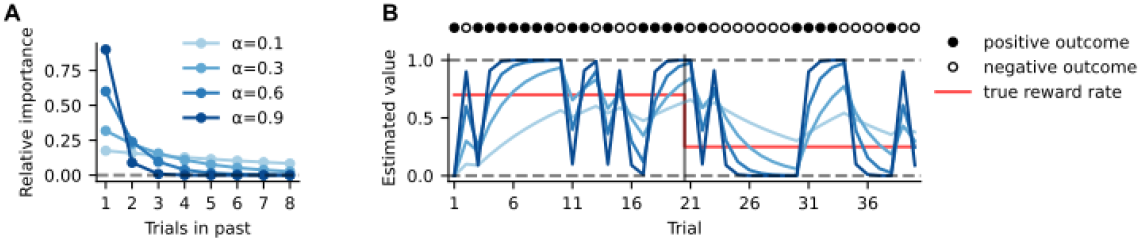
Delta learning rule. **A**, The delta learning rule, parameterized by the *α* learning rate, gives a recency-weighted average of past trials, where the relative importance (or discounting) of past outcomes decreases with growing learning rate parameter. Larger alpha puts more weight on recent outcomes, whereas alpha close to zero approximates the true mean of all the past outcomes (Sutton, 2018). **B**, Example session with stochastic outcomes. The true reward probability drops from 0.7 (first half) to 0.25 (second half, red step line). Filled and empty markers above the panel indicate rewarded and unrewarded trials, respectively. Higher alpha adapts quickly but steadily over-estimates the true reward rate (overfitting noise), whereas lower learning rate adapts more slowly but converges to the true reward rate more reliably.

**Figure S2.**
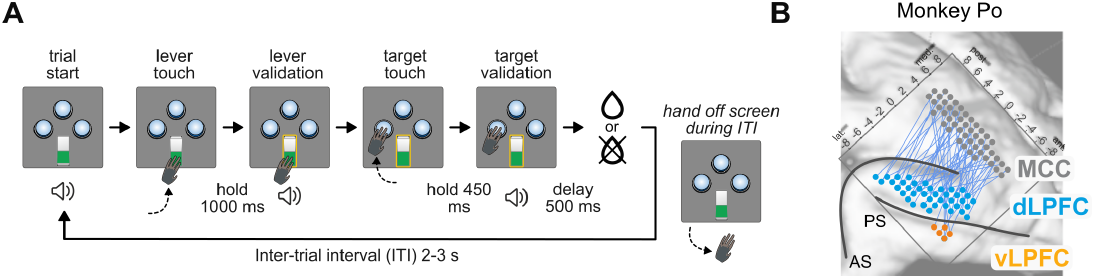
Experimental task and recording locations. **A**, Monkeys interacted with a three-armed bandit task through a touchscreen, where their actions produced visual feedback. Objects appeared on the screen as physical objects: the lever was represented as a binary flip switch, and the three options appeared as buttons that changed to a pressed state when touched. Unless validated, releasing these objects returned them to the original unpressed state. When the lever was validated, a yellow rectangle appeared around it and remained locked in the pressed position. Validation of both the lever and the target was indicated by a sound signal. Monkeys had to release the screen after the feedback to initiate a new trial. **B**, Recording sites for monkey PO (see KA in **Fig. 3A**).

**Figure S3.**
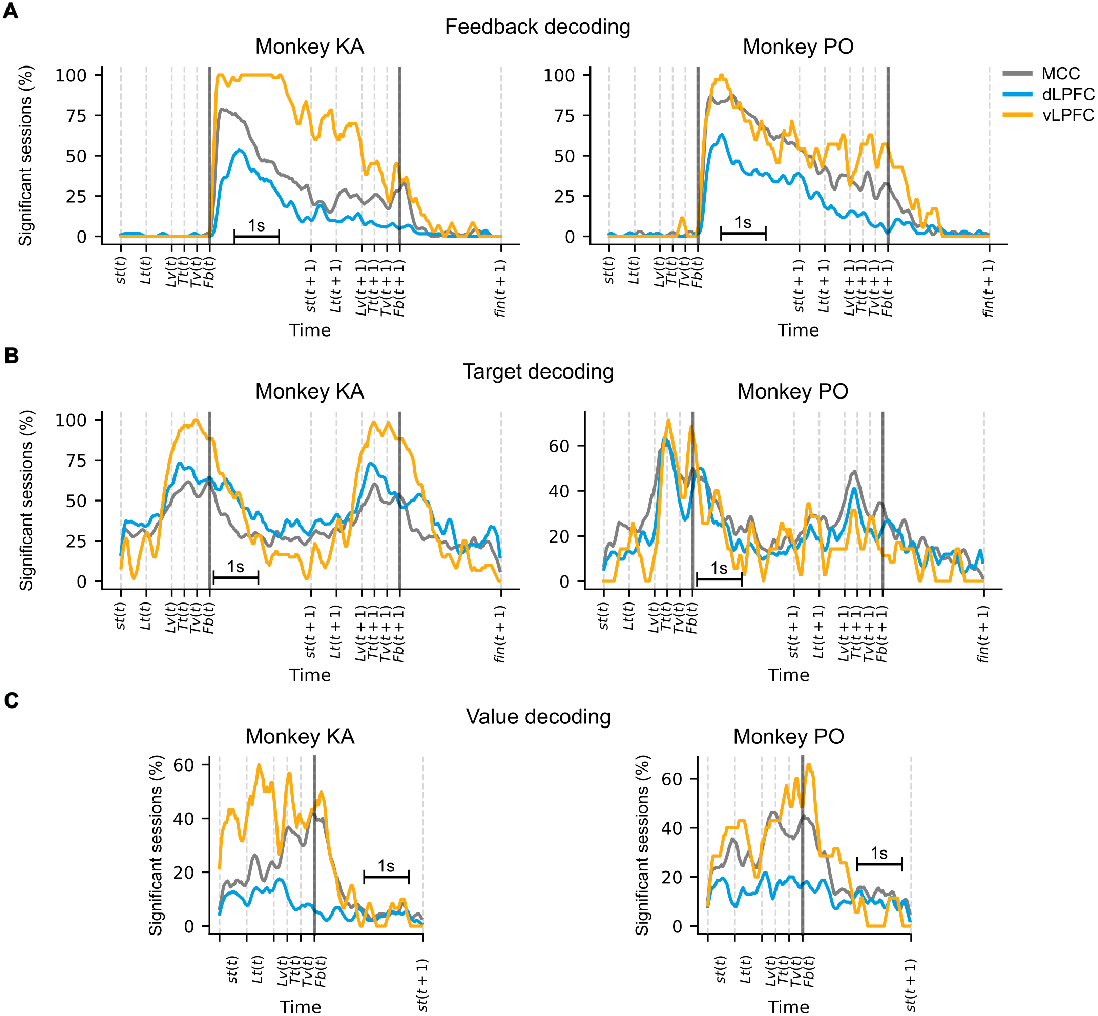
Per-monkey decoding performance of feedback, target and value. **A, B**, Feedback (A) and target (B) of trial *t*, decoded in the same and the subsequent trial. Both show strong decodability in the inter-trial interval and during the next trial. **C**, Value decoding in MCC and vLPFC ramps up towards the feedback event and declines sharply afterward.

**Figure S4.**
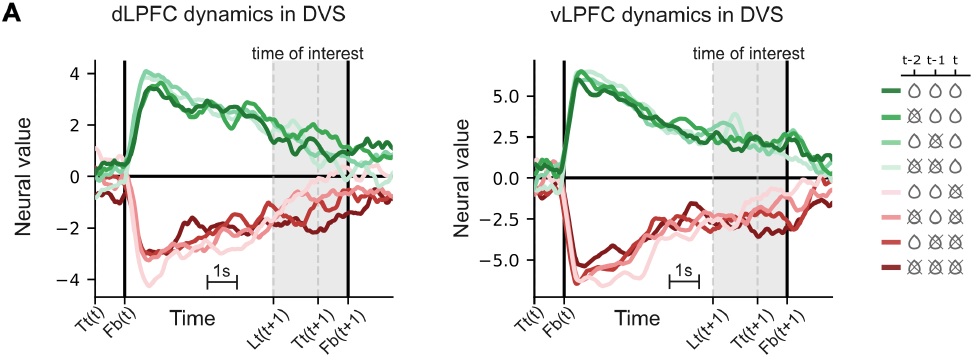
Dependence of LPFC subregions’ activity on outcome history. Activity in the decision-vector subspace (DVS; detailed in main text) is modulated by historical outcomes, especially during the task-engagement period (shaded background). Activity of composite-sessions are shown; see MCC in **Fig. 4B**

**Figure S5.**
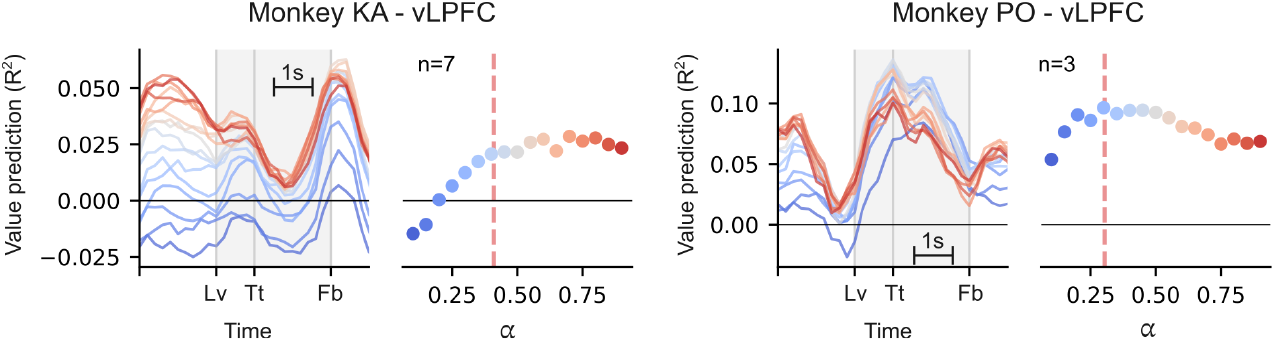
Behavior-derived value predicts neural activity under different learning rate parameters in vLPFC. Left panel: Correlation of calculated behavioral value with population activity as a function of varying learning rates of the foraging model. Shaded area: period of task engagement. Right panel: mean predictive strength of behavioral value as a function of learning rate, averaged across the task engagement period. Dashed line: learning rate from behavioral fit.

**Figure S6.**
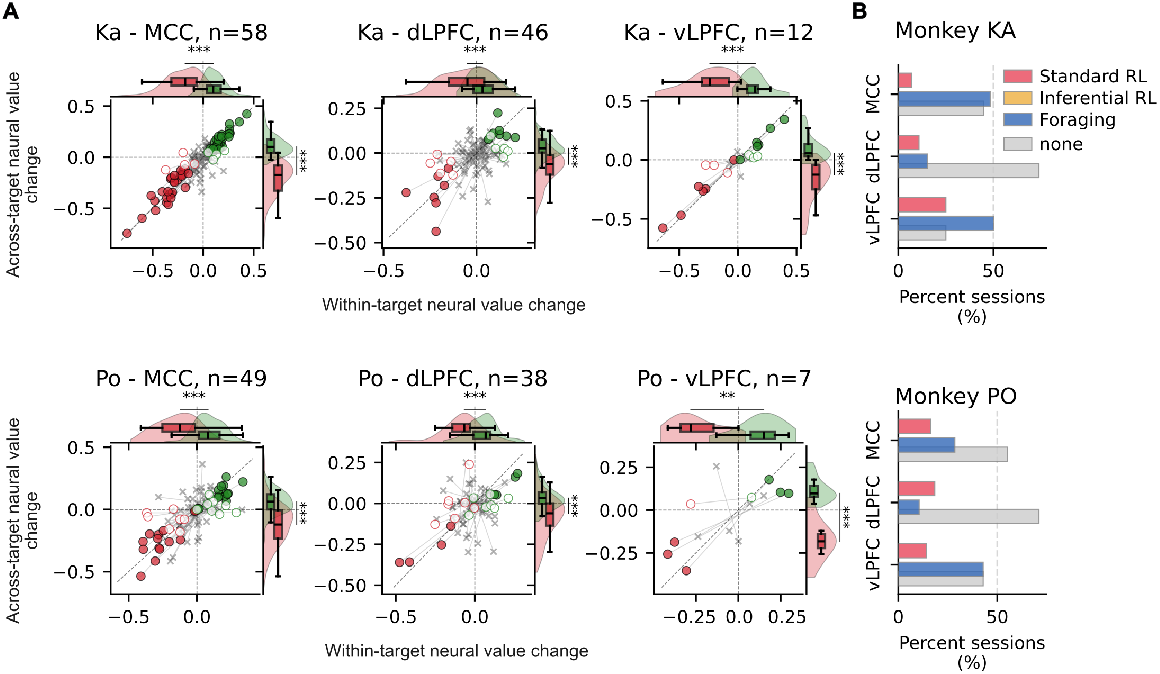
Per-monkey results of neural markers of foraging strategy. Recordings from both animals consistently support the foraging account of decision making.

## Supplementary Tables

**Table S1.** Statistical comparison of model performances. When fit to maximize reward, all models produced significantly different performances. We run 100 simulations of 100,000 trials per model, measure the performance in term of obtained reward rate, and compare this measure by pairwise Mann–Whitney U tests (two-sided; *n*_1_ = *n*_2_ = 100).

| Comparison | *U* | *p* |
| --- | --- | --- |
| Foraging vs. Inferential RL | 94 | $4.14 \times 10^{-33}$ |
| Foraging vs. Normative | 0 | $2.56 \times 10^{-34}$ |
| Inferential RL vs. Normative | 50 | $1.12 \times 10^{-33}$ |
| Foraging vs. Standard RL | 0 | $2.56 \times 10^{-34}$ |
| Inferential RL vs. Standard RL | 0 | $2.56 \times 10^{-34}$ |
| Foraging vs. WSLS | 0 | $2.56 \times 10^{-34}$ |
| Inferential RL vs. WSLS | 0 | $2.56 \times 10^{-34}$ |

